# The oncogenic fusion protein TAZ-CAMTA1 promotes genomic instability and senescence through hypertranscription

**DOI:** 10.1101/2022.12.01.518701

**Authors:** Emily Neil, Brian Rubin, Valerie Kouskoff

## Abstract

TAZ-CAMTA1 is a fusion protein found in over 90% of Epithelioid Hemangioendothelioma (EHE), a rare vascular sarcoma with an unpredictable disease course. To date, how TAZ-CAMTA1 initiates tumour formation remains unexplained. To study the oncogenic mechanism leading to EHE initiation, we developed a model system whereby TAZ-CAMTA1 expression is induced by doxycycline in primary endothelial cells. Using this model, we establish that upon TAZ-CAMTA1 expression endothelial cells rapidly enter a hypertranscription state, triggering considerable DNA damage. As a result, TC-expressing cells become trapped in S phase. Additionally, TAZ-CAMTA1-expressing endothelial cells have impaired homologous recombination, as shown by reduced BRCA1 and RAD51 foci formation. Consequently, the DNA damage remains unrepaired and TAZ-CAMTA1-expressing cells enter senescence. Knockout of *Cdkn2a*, the most common secondary mutation found in EHE, allows senescence bypass and uncontrolled growth. Together, this provides a mechanistic explanation for the clinical course of EHE and offers novel insight into therapeutic options.

## Introduction

Epithelioid Hemangioendothelioma (EHE) is a rare sarcoma of vascular endothelial cells, with an unpredictable disease course ^1^. EHE is a heterogeneous disease, with tumours reported at multiple anatomic sites, however the lungs, liver and bones are most commonly affected ^2, 3^. Two mutually exclusive chromosomal translocations define EHE tumours, each involving one of the transcription co-factors TAZ and YAP1 ^4-6^. TAZ and YAP1 are downstream effectors of the Hippo pathway, and primarily initiate transcription by interacting with TEAD transcription factors. They are frequently activated in cancer, however rarely directly mutated ^4^. TAZ-CAMTA1 (TC), resulting from a t(1;3) translocation, is present in 90% of EHE tumours ^5, 7^. The remaining 10% of EHE cases harbour a t(X;11) translocation, resulting in the YAP1-TFE3 (YT) fusion protein ^6^. It is thought that TC and YT represent the initiating events in EHE, however secondary mutations are present in at least 55% of cases and are associated with more aggressive disease ^8, 9^.

TC contains the N-terminus of TAZ fused to the C-terminus of CAMTA1 ^5^. TC retains the TEAD binding domain and the S89 residue critical to the regulation of wild-type TAZ activity. When the Hippo pathway is active, S89 is phosphorylated by LATS1/2 leading to the inactivation of TAZ ^5, 10^. CAMTA1 is a calmodulin binding transcription factor, however its DNA binding domain is not present in TC ^5, 11^. Initial studies have shown TC to be less sensitive to negative regulation by the Hippo pathway, despite S89 phosphorylation ^5, 12^. This is due to its increased nuclear localisation, driven by the nuclear localisation signal (NLS) contributed by CAMTA1 ^12^. Due to its interaction with TEADs, the transcriptional programme of TC bares similarities with the canonical TAZ signature ^5, 9, 13^. The CAMTA1 moiety contributes by recruiting the ATAC complex and increasing chromatin accessibility ^14^. This results in the EHE transcriptome being distinct from other endothelial neoplasms, but being conserved between EHE mouse models and human disease ^9^.

Genomic instability is a hallmark of cancer, whereby increased DNA damage allows mutations to accumulate ^15^. In cancer, DNA damage can arise from multiple sources, including replication stress ^16^. Hypertranscription upon oncogene activation is an increasingly recognised mechanism behind this, as replication forks collide with the transcriptional machinery and R-loops ^16, 17^. Replication induced double strand breaks (DSBs) result in S phase arrest whilst repair takes place ^18, 19^. Homologous recombination (HR) is a DSB repair pathway which dominates during S phase, and is typically error-free ^20^. HR involves resection of DSB ends by nucleases to create ssDNA overhangs, on which RAD51 is loaded by the BRCA2-PALB2 complex ^21^. The RAD51 nucleofilament can then invade the sister chromatid which is used as a template for accurate repair ^20, 22^. BRCA1 is a key HR effector, forming distinct complexes which function at multiple stages in the pathway ^23^. BRCA1 binding partners include Abraxas and RAP80 for recruitment to DSBs, CtIP for end resection, and BRCA2-PALB2 for RAD51 deposition ^21, 24, 25^. BRCA1 inactivation in cancer leads to genomic instability as HR is impaired^26^.

HR deficiency in cancer can result in oncogene induced senescence (OIS) ^27^. This tumour suppressive mechanism occurs when DNA damage is unable to be repaired, causing cells to undergo permanent growth arrest ^28^. Senescent cells are characterised by p16 expression, distinct morphology, and the secretion of inflammatory cytokines, termed the senescence-associated secretory phenotype (SASP) ^29^. The SASP recruits macrophages to clear senescent cells, however also contributes to an inflammatory, oncogenic microenvironment when senescent cells accumulate in a tumour ^30, 31^.

While transgenic mouse models have demonstrated the oncogenic potential of TC, we are still lacking cellular model systems to dissect the initiating molecular events driven by TC to promote EHE tumorigenesis. Here, we describe a model for generating primary endothelial cells from mouse embryonic stem cells (mESCs) harbouring a doxycycline inducible TC expression system. Upon TC induction, endothelial cells rapidly acquire transcriptomic and phenotypic features characteristic of human EHE. Using this model system to investigate how TC controls tumour initiation, we uncover a critical role for TC in promoting genomic instability via hypertranscription and DNA damage response interference.

## Results

### TC expression induces cell cycle arrest in primary endothelial cells

As wild-type TAZ is known to regulate S phase entry and proliferation in endothelial cells ^32^, we first determined if TC expression had a similar effect. The TIE2^+^FLK1^+^ endothelial progenitor cell population was isolated from differentiating mESCs and further cultured in endothelial promoting culture conditions (Supplementary Fig. 1a-d). The resulting cells expressed endothelial markers (TIE2, FLK1, and low level of CD31 and VE-cadherin) and formed cord-like structures in Matrigel plugs (Supplementary Fig. 1c, d). Addition of dox induced the co-expression of FLAG-tagged TC and GFP via an IRES sequence, which enables the tracking of TC-expressing cells by GFP expression in both live and fixed cells (Supplementary Fig. 1e, f). To investigate TC-expression over time, dox was added to endothelial cells and flow cytometry analysis was carried. FLAG epitope staining revealed the presence of two distinct populations expressing low or high level of the TC protein (Fig 1a). At 48h hours post induction, there was a significantly lower frequency of TC high endothelial cells compared to TC low, despite initially being of similar proportions (Fig. 1b). The proportion of TC high cells continued to decrease throughout the time course. This was further evidenced by Ki67 expression, which marks actively dividing cells. TC high and low populations initially had high frequencies of Ki67^+^ cells at 24h post dox induction compared to TC^neg^ and no dox controls. The frequency of Ki67^+^ cells was reduced significantly by 96h within in TC high and TC low populations (Fig. 1c, d, Supplementary Fig. 1g). This suggests that TC-expressing endothelial cells were unable to maintain an initially high proliferation rate, leading to a reduction in the frequency of TC high cells over time.

**Figure 1.**
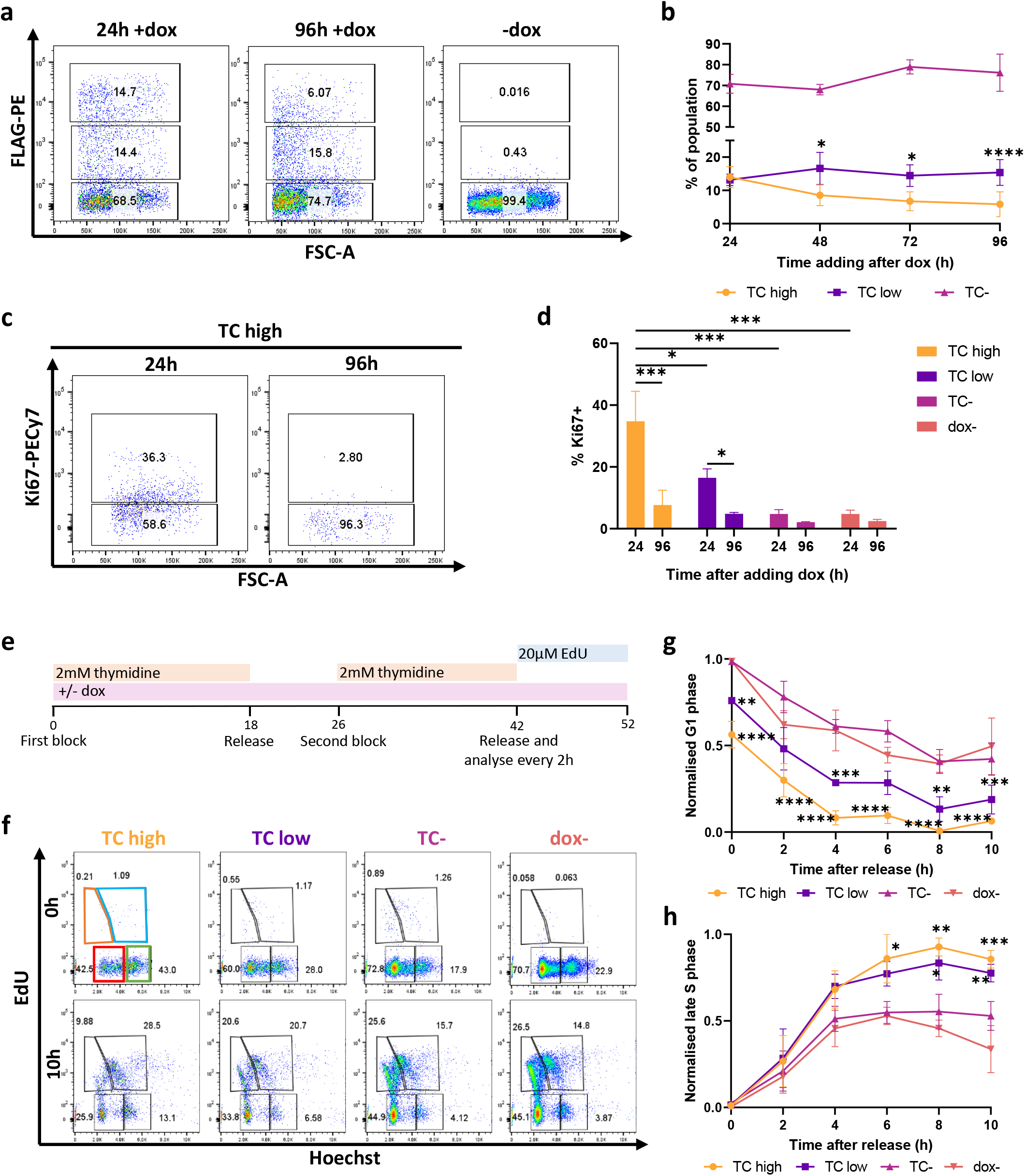
TC expression in endothelial cells initially results in cell cycle arrest. **a** Representative flow cytometry plots showing the frequency of TC expressing cells against FSC-A at 24h, and 96h with or without addition of doxycycline (dox) to induce TC expression. **b** The frequency of TC expressing endothelial cells at 24h, 48h, 72h and 96h after addition of doxycycline to induce TC expression, as measured by flow cytometry, n=4. Statistical significance was determined by two-way ANOVA with Tukey’s multiple comparisons test, n=3. **c** Representative flow cytometry plots showing the frequency of Ki67+ endothelial cells within the TC high population, at 24h and 96h after dox induction. **d** The frequency of Ki67+ endothelial cells at 24h and 96h after dox induction, within TC high, low, negative and uninduced populations, n=3. Statistical significance was determined by two-way ANOVA with Sidak’s multiple comparisons test. **e** Experimental timeline showing the double thymidine block protocol for cell cycle synchronisation at the G1/S phase boundary. **f** Representative flow cytometry plots showing EdU incorporation against Hoechst staining in TC high, TC low, TC- and uninduced endothelial cell populations at 0h and 10h after releasing cells from G1/S phase block. This reveals four populations EdU-Ho low (G1 phase; red gate), EdU+ Ho low (early S phase; orange gate), EdU+ Ho high (late S phase; blue gate), and EdU-Ho high (G2 phase; green gate). **g** Normalised frequency of cells in G1 phase over 10h following release from thymidine block, in TC high, TC low, TC- and uninduced endothelial cell populations, n=3. **h** as **g** but for normalised frequency of cells in late S phase, n=3. Statistical significance was determined by two-way ANOVA with Dunnett’s multiple comparisons test. In all panels error bars show SEM, and *p<0.05, **p<0.01, ***p<0.001, ****p<0.0001.

To investigate cell cycle progression in TC expressing endothelial cells, a double thymidine block was carried out to synchronise cells at the G1/S phase boundary (Fig. 1e). Thymidine was added for 18h to the endothelial cultures, followed by an 8h block-release, then an additional 16h thymidine block. At the end of the second block, cells were released and EdU was added to distinguish S phase cells. Hoechst staining was used to differentiate between G1 and G2 phase cells (Fig. 1e, f, Supplementary Fig. 1h). Flow cytometry revealed that TC-expressing populations had lower frequencies of G1 cells immediately after release compared to TC^neg^ and uninduced controls (Fig. 1g). Additionally, the proportion of TC-expressing cells in late S phase increased continually throughout the experiment, with a higher proportion than in the TC^neg^ populations by 8h (Fig. 1h). The proportion of TC^neg^ and uninduced endothelial cells in late S phase increased steadily, then began to decline (Fig. 1h). Together, these findings suggest that while initially TC expression caused increased proliferation, endothelial cells were subsequently arrested in late S phase, resulting in a lower proliferation rate compared to TC^neg^ and uninduced cells.

### TC expression causes DNA double strand breaks, independent of its interaction with TEAD

A well described mechanism behind S phase arrest is the accumulation of DNA damage ^18, 19^. Cells become arrested in S phase whilst damage is repaired to stop mutations being passed on to daughter cells. At sites of DSBs, γH2AX accumulates as H2AX is phosphorylated at S139 by the ATM/ATR kinases ^19, 33^. To address this potential cause for the S phase arrest observed in TC-expressing cells, endothelial cells were stained for γH2AX 24 hours after induction of TC expression, to assess the present of DSBs. This revealed that TC expression resulted in a substantial increase in γH2AX foci numbers, with a similar frequency of cells containing over 10 γH2AX foci to that of cells treated with H_2_O_2_ (Fig. 2a, b). An increased number of γH2AX^+^ cells was also observed upon induction of TC S51A expression, a mutant TC which cannot interact with TEAD factors ^34^ (Fig. 2a-b). γH2AX staining was already noticeable 4h after induction (Supplementary Fig. 2), suggesting that DSBs were a direct consequence of TC expression and independent of the interaction with TEAD factors. The number of foci in both dox and H_2_O_2_ treated populations was significantly higher than in uninduced endothelial cells at both time points (Fig. 2b, Supplementary Fig 2).

**Figure 2.**
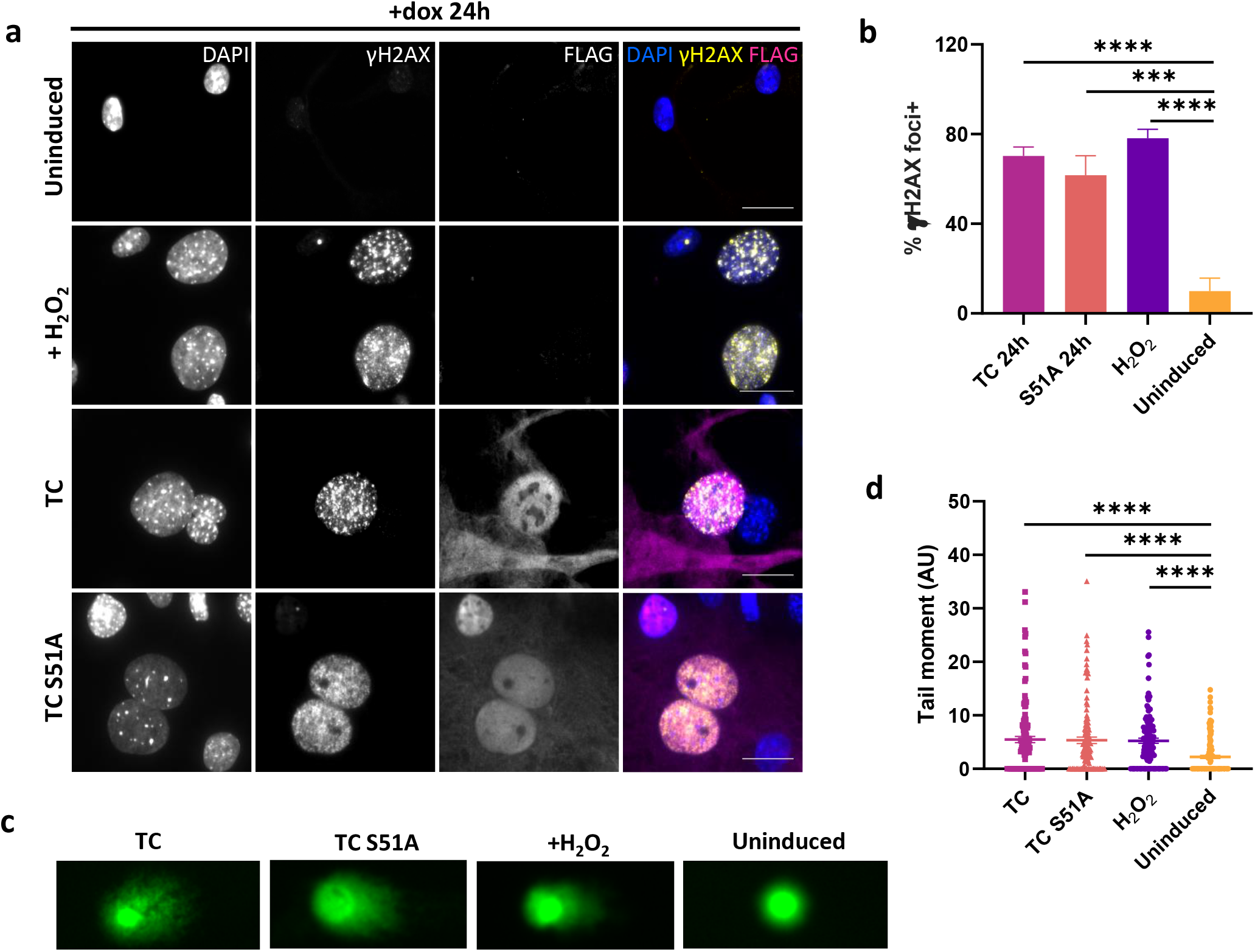
TC expression causes endothelial cells to accumulate DNA double strand breaks. **a** Representative images showing endothelial cells treated with dox for 24h to induce TC expression or not. Cells treated with 30μM H_2_O_2_ for 4h were used as a positive control. Cells were stained with DAPI (nuclei; blue), FLAG antibody (TC; magenta), and γH2AX antibody (phospho-H2AX; yellow). All imaging was performed on a Zeiss fluorescence widefield microscope using a 63x oil immersion objective. Scale bars=20μm. **b** Frequency of endothelial cells positive for γH2AX foci from imaging experiments as presented in **a**. Cells with more than 10 foci were considered positive. A minimum of 150 cells per condition were analysed, n=4. Significance was calculated by one-way ANOVA and Dunnett’s multiple comparisons test. **c** Representative images from neutral comet assay to detect double strand breaks in DNA. Day 10 endothelial cells were either left untreated, incubated with dox for 24h to induce TC or TC S51A expression, or treated with 30μM H_2_O_2_. Scale bars=50μm. **d** Graph showing the calculated Olive tail moment from the four conditions in c. A minimum of 125 cells per condition were analysed. Statistical significance was determined by one-way ANOVA with Sidak’s multiple comparisons test, n=3. In all panels error bars show SEM, and *p<0.05, **p<0.01, ***p<0.001, ****p<0.0001.

To independently confirm the presence of DSBs in TC-expressing cells, a neutral comet assay was performed which detects DNA fragmentation, indicative of DSBs, by single-cell gel electrophoresis (Fig. 2c, d). Untreated endothelial cells were compared to cells induced with dox for 24 hours or treated with H_2_O_2_ for 4 hours. The tail moment was then calculated to quantify DNA damage ^35^. Comet assay revealed that TC and TC S51A expressing cells, similar to H_2_O_2_ treated cells, had a significantly larger tail moment compared to untreated cells, of which most had no tail (Fig. 2c, d). These data revealed that endothelial cells acquire a large number of DSBs immediately upon TC expression, leading to cell cycle arrest. Moreover, this effect was independent of TC interaction with TEAD factors.

### TC expression mediates hypertranscription and replication stress in endothelial cells

Next, we sought to determine the mechanism behind TC induced DSBs. Oncogene activation can result in DNA damage via multiple mechanisms, however we investigated replication stress due to the short time period by which DSBs occur after TC expression ^16, 36^. We performed RNA-seq experiments 24 h after dox addition to compare the transcriptomes of TC high, low and negative endothelial cells to uninduced controls (Supplementary Fig. 3a, b). GSEA enrichment plots established that upon TC expression, primary endothelial cells derived from differentiating mESCs acquired a transcriptomic signature characteristic of EHE tumours and had a significant enrichment for TAZ/YAP targets ^9, 12^ (Fig. 3a, b, Supplementary Fig. 3c-d). Transcriptomic analysis also revealed a considerable number of differentially expressed genes (DEGs) in endothelial cells 24h after TC induction (Fig. 3c). This is suggestive of hypertranscription, a state whereby activity of RNA polymerases is increased. Hypertranscription is increasingly recognised as a source of replication stress in cancer cells, causing DSBs as a result of R-loop formation and increased replication-transcription conflicts ^37, 38^. Transcriptomic analysis revealed that expression of RNA polymerase II and III subunits were increased in TC expressing populations, associated with an enrichment for RNA metabolism gene set (Fig. 3d, e). To further investigate TC-induced hypertranscription, an RNA imaging assay was used to quantify nascent RNA (Fig. 3f, g). This revealed that, 24h after induction, TC-expressing cells had a higher amount of nascent RNA in their nuclei compared to uninduced controls (Fig. 3g). Moreover, immunofluorescence microscopy also revealed increased R-loop formation in dox treated endothelial cells (Fig. 3h, i). Together, this provides further evidence that hypertranscription occurs upon TC expression in endothelial cells, and that replication stress is the likely mechanism behind DSB formation and cell cycle arrest.

**Figure 3.**
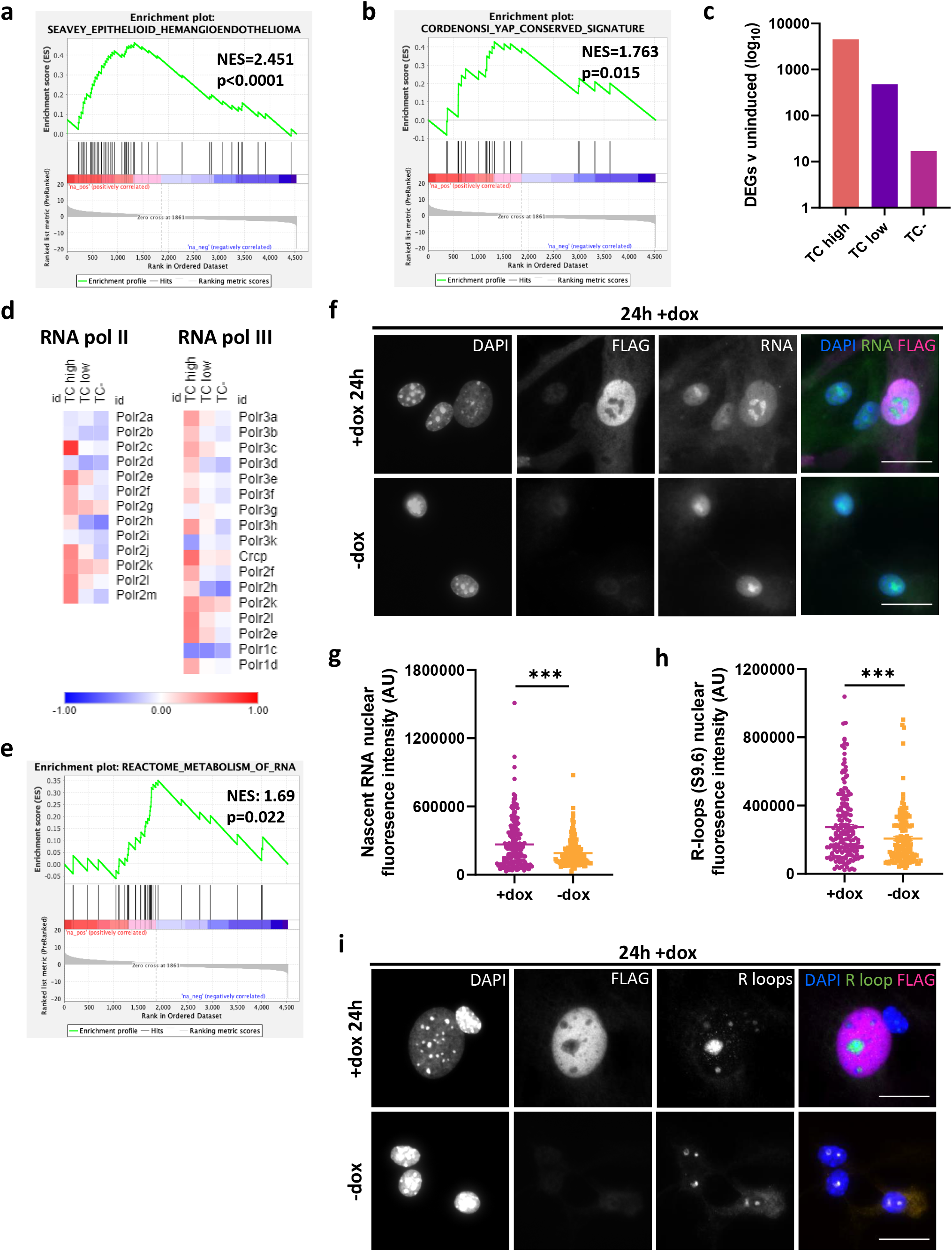
TC expression mediates hypertranscription and replication stress in endothelial cells. **a** GSEA enrichment plots showing significant positive enrichment for the Seavey Epithelioid Hemangio Endothelioma gene set in TC high endothelial cells. **b** GSEA enrichment plots showing significant positive enrichment for the Cordenonsi YAP conserved signature gene set in TC high endothelial cells. **c** The number of significantly differentially expressed genes (DEGs) determine from RNA-seq data in TC high, TC low and TC-populations when compared to uninduced controls. **d** Heatmaps showing log2 fold change in RNA expression of RNA polymerase II and III subunits when comparing TC high, TC low and TC-endothelial cell populations to uninduced controls. **e** GSEA enrichment plots showing significant positive enrichment for the reactome RNA metabolism gene set in TC high endothelial cells. **f** Representative images showing nascent RNA staining after endothelial cells were incubated with 5-EU for 1 hour. Cells were stained for 5-EU (nascent RNA; green), DAPI (nuclei; blue) and FLAG (TC; magenta). Imaging was performed on a Zeiss fluorescence widefield microscope using a 63x oil immersion objective. Scale bars=20*μ*m. **g** Fluorescence intensity of 5-EU staining as shown in **f**, to quantify nascent RNA. Statistical significance was determined with an unpaired t-test, n=3. **h** Fluorescence intensity of S9.6 antibody staining as shown in **i**, to quantify the presence of R-loops. Statistical significance was determined with an unpaired t-test, n=3. **i** Representative images showing S9.6 antibody staining to visualise R-loops. Cells were stained for R-loops (green), DAPI (nuclei; blue) and FLAG (TC; magenta). Imaging was performed on a Zeiss fluorescence widefield microscope using a 63x oil immersion objective. Scale bars=20*μ*m. In all imaging experiments, at least 150 cells per condition were analysed, n=3. In all panels error bars show SEM, and *p<0.05, **p<0.01, ***p<0.001, ****p<0.0001.

### Homologous recombination is impaired in TC expressing endothelial cells

To understand how DSBs generated upon TC expression might contribute to EHE tumorigenesis, we next investigated the ability of these cells to repair DNA damage. As TC causes S phase arrest, and HR is the dominant pathway in S phase, we investigated the activity of key HR proteins in TC-expressing endothelial cells. ^20^. Using immunofluorescent imaging, we visualised the expression and localisation of BRCA1 protein, which is recruited to DSBs and mediates the formation of DNA repair complexes ^23^. These experiments revealed that in TC-expressing endothelial cells, the number of BRCA1 foci+ cells was reduced compared to H_2_Otreated control populations, despite similar amounts of γH2AX foci (Fig. 4a, b). To further investigate, we next determined the amount of RAD51 foci in TC expressing endothelial cells. RAD51 is downstream in the HR pathway, and is recruited to the ssDNA overhangs produced after DSB end resection, where it mediates strand invasion ^20, 22^. RAD51 deposition is dependent on BRCA1 forming a complex with BRCA2 and PALB2 ^21^. Again, this revealed that in TC-expressing cells RAD51 foci+ cells were significantly less prevalent than in H_2_O_2_ treated controls (Fig. 4c, d). Together, these data suggest that the HR arm of DNA damage response is impaired in TC-expressing cells.

**Figure 4.**
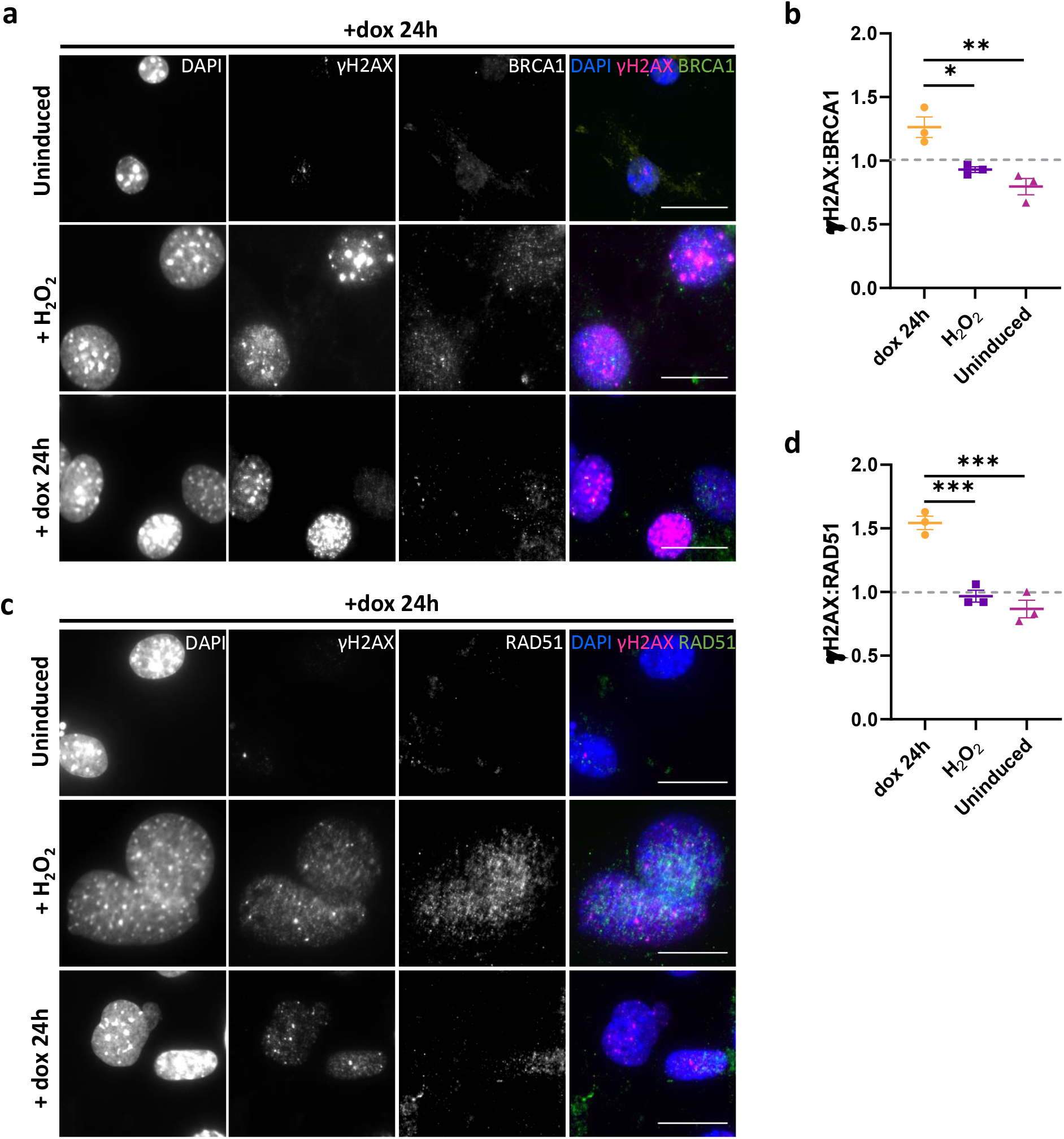
Homologous recombination is impaired in TC expressing endothelial cells. **a** Representative images showing endothelial cells that were left untreated, incubated with dox for 24h to induce TC expression, or treated with 30μM H_2_O_2_. Cells were stained with DAPI (nuclei; blue), γH2AX (magenta), and BRCA1 (green). **b** Ratio of γH2AX to BRCA1 foci positive endothelial cells from imaging experiments as shown in (A). **c** Representative images showing endothelial cells that were left untreated, incubated with dox for 24h to induce TC expression, or treated with 30μM H_2_O_2_. Cells were stained with DAPI (nuclei; blue), γH2AX (magenta), and RAD51 (green). **d** Ratio of γH2AX to RAD51 foci positive endothelial cells from imaging experiments as presented in **c**. All imaging was performed on a Zeiss fluorescence widefield microscope using a 63x oil immersion objective. Scale bars=20*μ*m. A minimum of 150 cells per condition were quantified per experiment, n=3. Statistical significance was calculated with a one-way ANOVA and Sidak’s multiple comparisons test. In all panels error bars show SEM, and *p<0.05, **p<0.01, ***p<0.001, ****p<0.0001.

### TC expression in endothelial cells results in oncogene-induced senescence and genomic instability

HR-deficient cancer cells may undergo OIS, as the amount of DSBs overwhelms compensatory repair pathways ^27^. Hence, we next investigated whether this was the case for TC-expressing endothelial cells in our model system. As senescent cells do not proliferate, this might explain the reduced cell growth observed for TC-expressing endothelial cells as shown above (Fig. 1). To explore this possibility, endothelial cells were exposed to dox for 24h to induce TC expression, then p16 expression was determined by immunofluorescence, as this protein is a known positive regulator of senescence (Fig. 5a, b). Importantly, *CDKN2A*, which encodes p16, is the most common secondary mutation in EHE, and these tumours are often more aggressive ^8^. Fluorescence microscopy revealed that dox induced TC-expressing endothelial cells had a significantly higher expression of p16 than untreated controls. Interestingly, some cells presented with a cytoplasmic expression of TC and a nuclear expression of p16, suggesting a transition where, before being shut down, TC is first translocated to the cytoplasm concomitant to p16 expression (Fig. 5a; white arrowhead).

**Figure 5.**
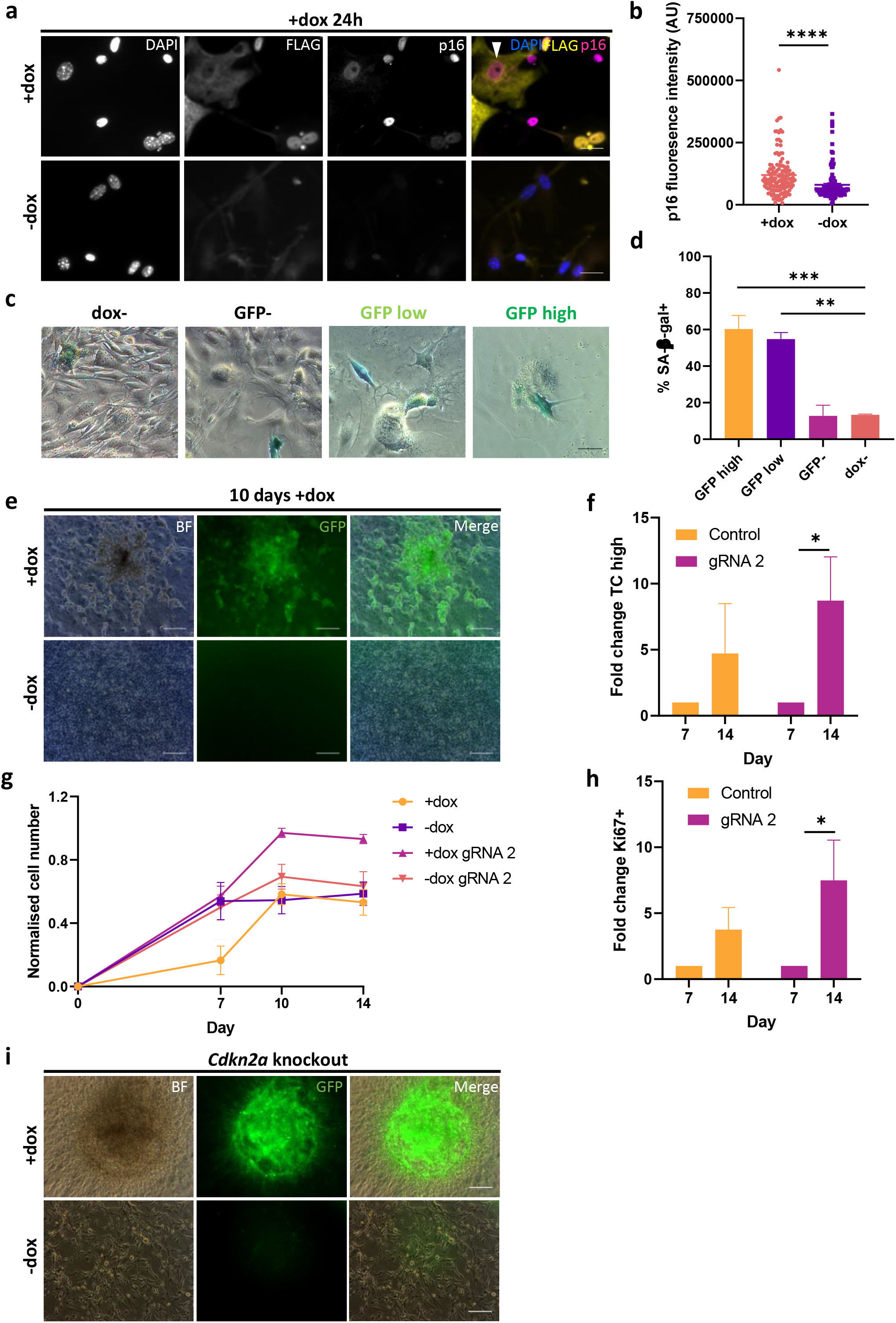
TC expression in endothelial cells results in oncogene-induced senescence, which can be overcome by a secondary mutation. **a** Representative images showing endothelial cells that were left untreated or incubated with dox for 24h to induce TC expression, then stained for DAPI (nuclei; blue), p16 (magenta), and FLAG (TC; yellow). Imaging was performed on a Zeiss fluorescence widefield microscope using a 40x objective. Scale bars=25*μ*m. **b** Fluorescence intensity of p16 staining as shown in a, to quantify its expression, n=3. Statistical significance was determined with an unpaired t-test. **c** Representative images of GFP (TC) high, low, negative and uninduced populations stained for SA-β-gal activity, 4 days after addition of dox. **d** Frequency of cells positive for SA-β-gal activity in GFP high, GFP low, GFP-, and dox-endothelial cell populations. A minimum of 150 cells per condition were analysed, n=3. Significance was calculated with a one-way ANOVA and Tukey’s multiple comparisons test. **e** Representative images showing the formation of GFP+ foci, whereby GFP is a marker for TC expressing endothelial cells, after 10 days exposure to dox. Scale bars=200*μ*m, n=3. **f** Fold change of TC high population between days 7 and 14 after nucleofection with control and *Cdkn2a* knockout guide RNA, n=4. Statistical significance was calculated with a two-way ANOVA and Sidak’s multiple comparisons test. g Normalised cell number in *Cdkn2a* knockout endothelial cells, treated with dox or not, over 14 days, n=4. Statistical significance was calculated with a two-way ANOVA and Sidak’s multiple comparisons test. All P-values are outlined in Supplementary Table 1. **h** Fold change of Ki67 expression within the TC high population between days 7 and 14 after addition of dox in comparison to uninduced endothelial cells, n=4. Statistical significance was calculated with a two-way ANOVA and Sidak’s multiple comparisons test. **i** Representative images showing the formation of GFP+ foci, whereby GFP is a marker for TC expressing endothelial cells, after *Cdkn2a* knockout. Scale bars=200*μ*m, n=4. In all panels error bars show SEM, and *p<0.05, **p<0.01, ***p<0.001, ****p<0.0001.

We also assessed senescence-associated β-galactosidase activity (SA-β-gal) in endothelial cells expressing TC or not, another hallmark of senescent cells ^29^. To this end, GFP sorted endothelial cells were cultured for 4 days and subjected to SA-β-gal staining (Fig. 5c, d). This revealed a higher proportion of cells with SA-β-gal activity in GFP high and low populations, compared to GFP^neg^ and untreated endothelial cells (Fig. 5d). Of note, the GFP high and low populations also took on the characteristic flat morphology of senescent cells, whereas GFP^neg^ and uninduced cells retained their endothelial morphology (Fig. 5c). Together, these data revealed that TC expression resulted in OIS in endothelial cells, following increased in p16 expression.

To study the long-term consequences of TC expression, endothelial cells were cultured for 4 weeks, with dox added every three days to maintain TC expression. After 10 days, small cell aggregates began to form, which were not present in dox-populations (Fig. 5e). These aggregates were GFP positive, suggesting sustained TC expression. Hence, we hypothesised that a subset of TC-expressing cells may bypass senescence by acquiring a secondary mutation. CDKN2A was shown to be the most common secondary mutation in EHE, conferring increased tumour aggressiveness ^8^. As a proof of principle of the additional effect of secondary mutation, we investigated the effect of CRISPR/Cas9 *Cdkn2a* knockout on endothelial cell growth, expressing TC or not (Fig 5f-i, Supplementary Fig. 4, 5). Here, flow cytometry revealed that the proportion of TC high cells increased significantly by day 14 in the *Cdkn2a* knockout condition, in contrast to the dox+ controls (Fig. 5f, Supplementary Fig. 4). Additionally, *Cdkn2a* knockout in dox treated cells resulted in a significant increase in cell number compared to control dox treated cells at all the time points tested (Fig. 5g, Supplementary Fig. 5, Supplementary Table 1) and the formation of very large GFP^+^ cell clusters (Fig. 5i). *Cdkn2a* knockout had little effect on untreated populations. Moreover, the number of Ki67+ TC high cells increased in all conditions between days 7 and 14, however this was only significant in *Cdkn2a* knockout populations (Fig. 5h, Supplementary Fig. 5). This provides evidence that a secondary, senescence-bypassing mutation is able to overcome the growth arrest imposed by TC expression.

### The interaction between TC and TEAD is required to sustain proliferation

Using a TC-S51A mutant, we showed above that the interaction between TC and TEAD was not essential for the generation of DSBs upon TC initial expression in endothelial cells (Fig. 2, Supplementary Fig. 2). However, many previous studies have highlighted that the interaction of TC with TEAD is important for tumorigenesis ^9, 12, 13^. Therefore, we aimed to determine if the TC-TEAD interaction had a role in maintaining tumorigenesis in endothelial cells upon longer-term culture. Here, we generated endothelial cells and added dox to induced TC-S51A expression. Over the following 14 days, we analysed cell number and TC-S51A expression in comparison to untreated cells. Unlike TC-expressing endothelial cells, we did not observe GFP^+^ colony formation in TC-S51A expressing cultures after 10 days of dox induction (Fig. 6a). Moreover, TC-S51A expressing cultures remained at a lower cell number than untreated controls at all time points examined (Fig. 6b). This is in contrast to TC-expressing endothelial cells, which resulted in a comparable cell number to uninduced controls by day 14 (Fig. 5f-h). Similar to TC, the frequency of cells expressing TC-S51A did increase throughout the experiment (Fig. 6c, d). However, the number of FLAG high Ki67+ cells remained the same between days 7 and 14 (Fig. 6e, f). Overall, this shows that while the TC-TEAD interaction appears to be dispensable for DSB accumulation, it is required to sustain proliferation.

**Figure 6.**
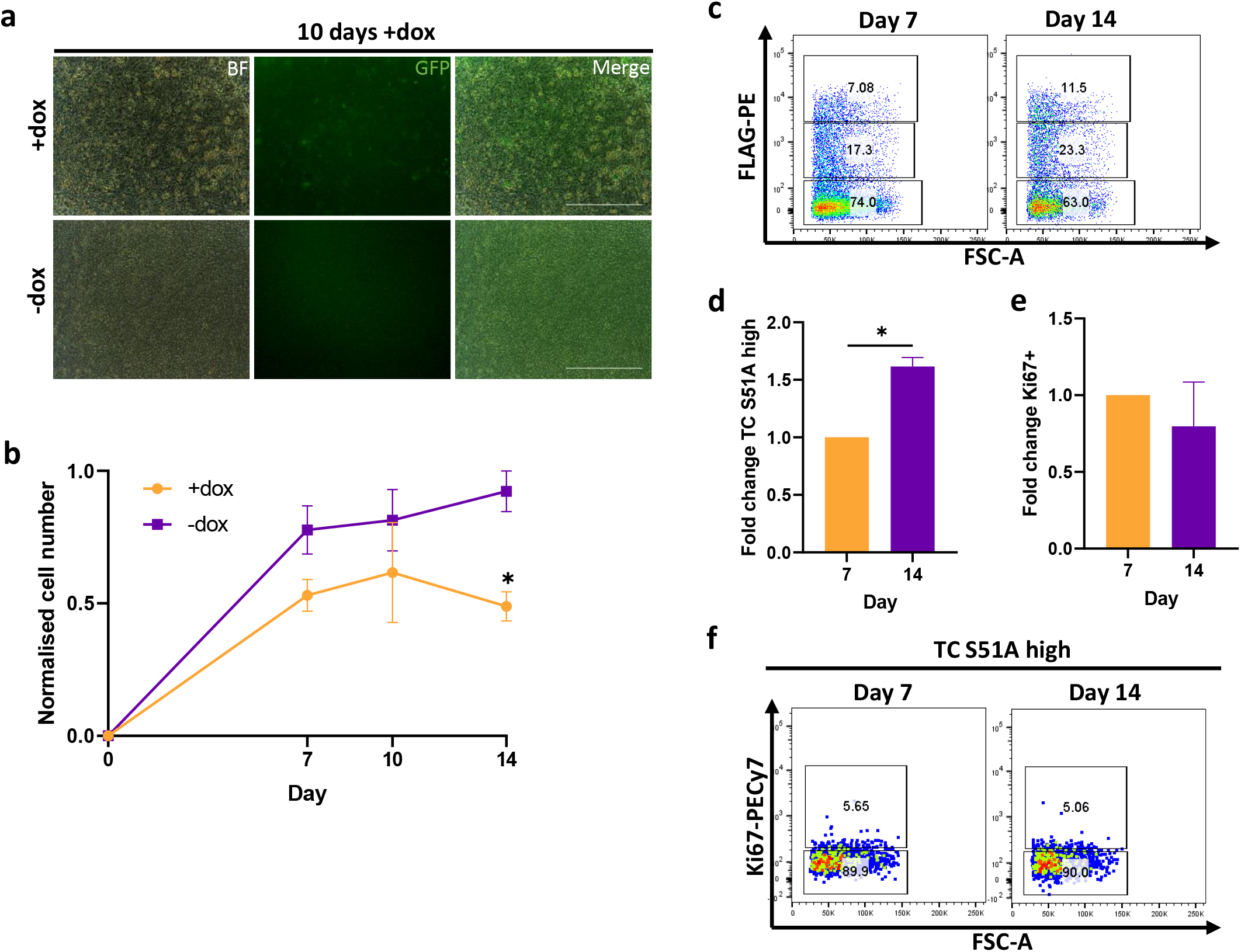
The interaction of TC and TEAD is required for maintenance of tumorigenesis. **a** Representative images showing endothelial cells treated with dox or not to induce TC S51A expression, whereby GFP is a marker for TC S51A, after 10 days exposure to dox. Scale bars=200*μ*m, n=3. **b** Normalised cell number in endothelial cells treated with dox or not to induce TC S51A expression, over 14 days, n=3. Statistical significance was calculated with a two-way ANOVA and Sidak’s multiple comparisons test. **c** Representative flow cytometry plots showing TC S51A expression against FSC-A at day 7 and 14. **d** Fold change of TC S51A high population between days 7 and 14 after addition of dox in comparison to uninduced endothelial cells, n=3. Statistical significance was calculated with a paired t-test. **e** Fold change of Ki67 expression within the TC S51A high population between days 7 and 14 after addition of dox in comparison to uninduced endothelial cells, n=3. Statistical significance was calculated with a paired t-test. **f** Representative flow cytometry plots showing Ki67 expression within the TC S51A high population against FSC-A at day 7 and 14. In all panels error bars show SEM, and *p<0.05, **p<0.01, ***p<0.001, ****p<0.0001.

## Discussion

Here, we describe an *in vitro* model for studying the consequences of TC expression in primary endothelial cells, and how this contributes to EHE development. Using this model, we established that hypertranscription upon TC expression generates DSBs and causes cell cycle arrest (Fig. 7). Impairment of the HR pathway leaves these DSBs unresolved, leading to OIS in most cases. However, this also leaves TC-expressing endothelial cells susceptible to acquiring a growth-promoting secondary mutation. Previous studies have highlighted TC expression as being the initiating event in EHE development, however have used either contextually non-relevant cell lines or mouse models which take a long time to generate tumours ^5, 9, 12^-^14, 39^. Using our model system, we provide a mechanistic explanation for previously unanswered questions about the unique clinical features of EHE within a contextually relevant, *in vitro* model.

**Figure 7.**
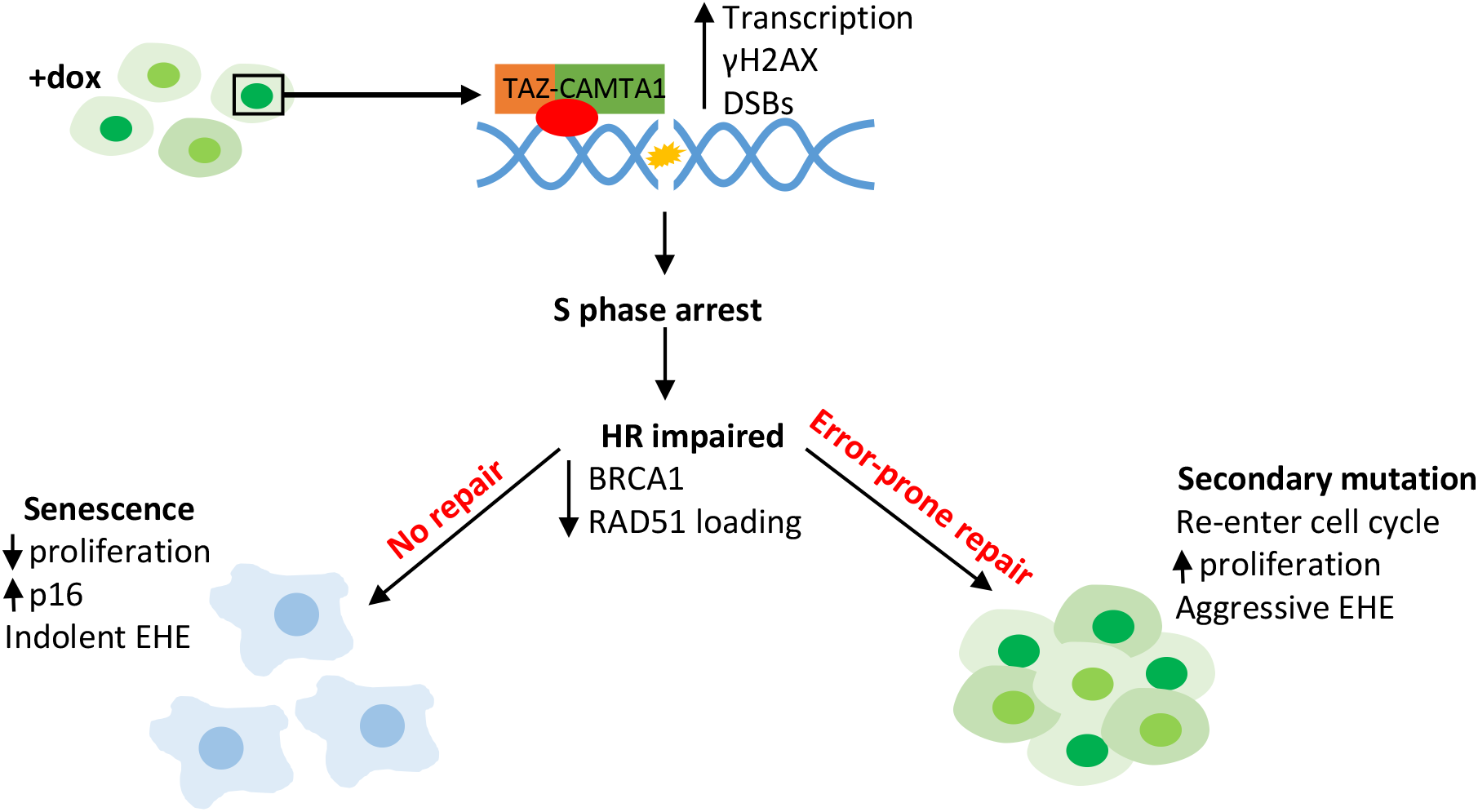
TC expression in endothelial cells results in senescence and genomic instability. Schematic showing how TC expression in endothelial cells results in hypertranscription, leading to the accumulation of DNA damage which in turn causes S phase arrest. As TC-expressing endothelial cells have impaired HR, as shown by reduced BRCA1 and RAD51 foci formation, many become senescent as the large amount of DNA damage is not repaired. These cells have increased p16 expression and remain arrested. In a subset of cells an error-prone DNA repair pathway will take over, leaving these vulnerable to acquiring growth-promoting secondary mutation.

EHE is described as having an unpredictable disease course; indolent tumours can switch to a highly aggressive phenotype ^1, 2^. We were able to replicate and provide mechanistic explanation for these clinical features. We show that upon TC expression, endothelial cells accumulate overwhelming DNA damage, initially causing increased p16 expression and cell cycle arrest. This later leads to SA-β-gal activity and a decreased growth rate, which are all hallmarks of OIS ^29^. We therefore propose that the indolent phase of EHE is likely due to the onset of OIS in TC expressing endothelial cells. OIS also explains the low mutational burden of EHE tumours in the face of genomic instability, as growth arrest is imposed before any error-prone repair mechanism can take place and mutations passed on to daughter cells.

We also reveal that in TC-expressing endothelial cells, the combination of replication-stress induced DNA damage and impaired HR contribute to genomic instability, a hallmark of cancer ^15^. Moreover, secondary mutations are present in at least 55% of EHE tumours, and these cases tend to be more aggressive ^8^. The most common of these secondary mutations affects *CDKN2A*, which encodes p16, a key positive regulator of senescence ^8, 29^. We were able to establish that *Cdkn2a* knockout in our model system allowed increased proliferation compared to *Cdkn2a* competent counterparts. This highlights the acquisition of a senescence-bypassing mutation as a mechanism behind the development of aggressive EHE.

Our model provides evidence that TC expressing endothelial cells have compromised DNA repair, specifically the HR pathway. This phenomenon is present in many cancers, with well-known examples including breast and ovarian tumours ^40^. In these tumours, error-prone pathways take over, leading to accumulation of mutations ^26^. Importantly, the discovery that HR is impaired in TC-expressing endothelial cells provides insight into potentially useful treatment options. Our data suggests that drugs targeting DNA repair, such as PARP or ATR inhibitors, may induce apoptosis preferentially in EHE tumour cells through synthetic lethality ^41, 42^.

Interestingly, our data highlights the striking similarities between the mechanism of action of TC and other fusion proteins found in sarcomas. Often, these fusion proteins dysregulate transcription by recruiting chromatin modifying complexes, as is seen for TC in EHE ^14, 43^. One of the best studied fusion protein is EWS-FLI1, which is found in Ewing sarcoma. Here, EWS-FLI1 expression induces hypertranscription and DNA damage, leading to growth arrest ^38, 44^. Another commonality between TC and EWS-FLI1 is the induction of HR deficiency, while the tumours maintain a low mutational burden ^38, 43, 45^. This is observed in Ewing sarcoma, but also in rhabdomyosarcoma (driven by PAX3/7 fusion proteins) and synovial sarcoma (SS18-SSX) ^46, 47^. Frequently, these fusion protein-defined sarcomas exhibit a loss of p16 which is associated with increased aggressiveness, suggesting a role for OIS ^44, 46, 48-50^. Together, these findings could provide insight into broadly relevant treatment strategies for EHE. Examples include targeting chromatin modifiers, CDK4/6 inhibition and senolytics, which have proven efficacious in the other sarcomas mentioned above ^43, 51, 52^. This is particularly important for rare cancers such as EHE, where studies are limited by sample size.

It would also be of interest to investigate the mechanisms behind HR-impairment in TC-expressing endothelial cells. Often, in other cancers, this is due to an inactivating mutation in a key HR protein, such as BRCA1/2, and represents the initiating event ^26^. This cannot be the case in EHE as the DNA damage and growth arrest phenotype is evident immediately after TC expression, and there is no common HR inactivating mutation across tumour samples ^8^. We hypothesise that TC may interfere with HR at the protein level, for example by inhibiting BRCA1 repair complex formation. Similar phenomena have been observed in Ewing sarcoma, where BRCA1 remains associated with the transcriptional machinery despite DSBs, and with oncogenic RAS, which induces BRCA1 chromatin dissociation ^27, 38^. Further study will be required to investigate how TC interferes with HR repair mechanisms.

Previous studies have demonstrated the ability of TC to interact with TEAD as vital to its transforming ability ^9, 12, 13^. In contrast, we show that the induction of DSBs is independent of the TC-TEAD interaction. We propose that this is due to the replication stress induced by increased transcription being primarily mediated by the CAMTA1 moiety. Previous reports suggest that CAMTA1 can recruit the ATAC complex for chromatin modification, which would permit the large transcriptomic changes required for induction of replication stress ^14^. Despite this, our model does show significant enrichment of canonical TAZ/YAP gene sets upon TC expression, in agreement with current studies. It is also evident that the TC-TEAD interaction is still crucial to EHE progression, as in longer term cultures TC-S51A expressing endothelial cells were unable to overcome growth arrest. In many published studies, transformed cell lines were used to study the effects of TC and TC-S51A ^12-14^. We propose that using already transformed cell lines mimics the effect of a secondary mutation, allowing tumorigenesis as soon as TC is expressed.

To summarise, we present here a model for generating endothelial cells from mESCs, which can be used to study EHE by inducing TC expression. This inducible *in vitro* model is very powerful as it allows generating large quantity of cells for analysing the immediate consequences of TC expression. Our model reveals that TC expression rapidly results in OIS and genomic instability, and that these processes may explain the clinical features of EHE. Understanding the molecular mechanism by which the oncogenic fusion protein TC promotes tumour progression will help devise novel treatments for EHE patients.

## Methods

### Maintenance of mouse embryonic stem cells

The mESCs contain an inserted construct into the HPRT locus whereby TC or TC S51A expression is induced upon the addition of doxycycline (dox) as previously described ^53^. Both TC and TC S51A sequences contain an N-terminal double FLAG epitope, and is followed by an IRES-GFP reporter cassette. In all experiments that required TC or TC S51A protein expression, dox was added to the culture media at a final concentration of 1 *μ*g/ml. Parallel control experiments were set without dox (uninduced). mESCs were expanded and maintained in DMEM-ES medium. This comprised of Dulbecco’s Modified Eagle Medium (DMEM; Sigma) supplemented with 15% foetal calf serum (FCS), 2 mM L-glutamine (Gibco), 50 U/ml penicillin-streptomycin (Sigma), 1% LIF conditioned media, and 0.15 mM Monothioglycerol (MTG; Sigma). Cells were maintained on a monolayer of Mitomycin C inactivated mouse embryonic fibroblasts (MEFs) until differentiation. mESCs were cultured at 37°C with 5 % COin a humidified incubator.

### Generation and maintenance of mESC-derived endothelial cells

To initiate the endothelial differentiation process, mESCs were passaged twice onto 0.1% gelatine coated plates without MEFs. The cells were first passaged in DMEM-ES, then in IMDM-ES. IMDM-ES contains IMDM supplemented with 15% FCS, 2 mM L-glutamine, 50 U/ml penicillin-streptomycin, 1% LIF conditioned media, and 0.15 mM MTG. Next EBs were generated by harvesting mESCs and transferring IMDM supplemented with 15% FCS, 2 mM L-glutamine, 50 U/ml penicillin-streptomycin, 0.5 ng/ml ascorbic acid (Sigma), 180 µg/ml transferrin (Sigma), 0.45 mM MTG and 5 ng/ml human vascular endothelial growth factor (hVEGF; Peprotech). EBs were cultured in non-tissue culture treated 10 cm dishes. From this point on, cells were maintained in low oxygen (5 % O_2_) conditions, at 37°C with 5% CO_2_ in a humidified incubator. At day 5 EBs were subject to cell sorting to isolate the TIE2+ FLK1+ population. After sorting, cells were cultured in endothelial cell medium, which contains IMDM with 10% FCS, 2 mM L-glutamine, 50 U/ml penicillin-streptomycin, 180 µg/ml transferrin, 0.45 mM MTG, 0.25 ng/ml ascorbic acid, 15% D4T conditioned medium, 5 ng/ml murine basic fibroblast growth factor (bFGF; Peprotech) and 5 ng/ml hVEGF. Cells were maintained on Matrigel (Corning) diluted to a protein concentration of 5.5 mg/ml in IMDM. Endothelial cell medium was replaced every 2-3 days and supplemented with dox as appropriate.

### Flow cytometry

For analysis of extracellular markers, cells were dissociated with TrypLE Express (Gibco) at the indicated time points. Cells were washed using FACS wash (1X PBS 10% FCS) to create single cell suspensions. Staining was carried out for 30 minutes at 4°C, using the antibodies listed in Supplementary Table 2. Cells were also analysed for GFP as a marker for TC or TC S51A expression, depending on the cell line used. After staining, cells were washed and resuspended in FACS wash prior to analysis. For intracellular analysis, cell pellets were resuspended in 2% paraformaldehyde (PFA; Alfa Aesar) in 1X PBS for 20 minutes at room temperature. Cell were then permeabilized using 0.1% saponin in FACS wash for 15 minutes at room temperature. Fluorophore pre-conjugated or unconjugated primary antibodies were then added to samples at the appropriate concentrations, then left to stain for 30 minutes at 4°C. If used, fluorophore pre-conjugated secondary antibodies were then added and incubated for 30 minutes at 4°C. Afterwards, cell samples were washed then resuspended in 0.1% saponin in FACS wash for analysis. All analysis was performed using an LSR Fortessa cytometer (Becton Dickinson).

### Cell sorting

For sorting the TIE2^+^FLK1^+^ cell population, EBs were harvested and allowed to settle under gravity, before dissociation using TrypLE Express. Cells were then resuspended in IMDM 10% FCS and passed through a 50 μm Filcon (Becton Dickinson), prior to staining with TIE2 and FLK1 antibodies (Supplementary Table 2), and Hoechst 33258 for 1 hour at 4°C. Cell samples were vortexed after 30 minutes to prevent clumping. After staining, samples were washed and resuspended in IMDM 10% FCS, and sorted using the FACS Aria or Influx cell sorters (Becton Dickinson) at 4°C. After sorting, TIE2^+^FLK1^+^ cells were collected and re-plated in conditions for endothelial cell culture, as described above.

### Cell cycle analysis

For cell cycle analysis, cells were synchronised at the G1/S phase boundary using a double thymidine block. Here, endothelial cells were treated with 2 mM thymidine (Sigma) in cell culture grade H_2_O_2_ for 18h then released into fresh media for 8 hours. Endothelial cells were then subject to a second block using 2 mM thymidine again for 16 hours, then released into fresh media. At this point, 20 *μ*M 5-ethynyl-2’-deoxyuridine (EdU) in DMSO was added, then samples were harvested every 2 hours for 10 hours in total. The cells were then fixed in 4% paraformaldehyde for 15 minutes and stored in 1X PBS overnight at 4°C. The next day, staining and the click chemistry reaction to detect EdU incorporation was carried out using the Click-iT Plus EdU Alexa Fluor 647 Flow Cytometry Assay Kit (Invitrogen), as per manufacturer’s instructions. Cells were then stained with Hoechst 33258 to analyse DNA content, and an anti-FLAG tag antibody (Supplementary Table 2) to determine TC expression level.

### RNA sequencing and analysis

To collect samples for RNA sequencing, mESC-derived endothelial cells were cultured for 10 days after TIE2^+^ FLK1^+^ antibody sorting, before inducing TC expression or not with dox. 24 hours later, endothelial cells were harvested and sorted by flow cytometry into four populations; TC high, TC low, TC neg, and uninduced. TC expression levels were defined by GFP expression (Supplementary fig. 1f, 3a). After sorting, cell pellets were resuspended in RLT lysis buffer from the RNeasy Mini Kit (Qiagen) and stored at −80°C until sufficient samples for three biological repeats had been collected. RNA extraction was then performed on all samples at the same time, using the RNeasy Mini Kit, as per manufacturer’s instructions. RNA concentration was quantified using the Qubit RNA BR Assay Kit (Invitrogen) and Qubit 3 fluorometer (Invitrogen). Sequencing was performed by the Genomic Technologies Core Facility at the University of Manchester, using the Illumina HiSeq 4000 system. Unmapped paired-end sequences were assessed by FastQC, then sequence adapters were removed and reads trimmed using Trimmomatic_0.36. The reads were then aligned to the mm10 reference mouse genome, and gene counts were calculated using annotation from GENCODE M25 using STAR_2.7.2b. Normalisation, principle component analysis, and differential gene expression was calculated in DESeq2_1.20.0 using default settings. This bioinformatics analysis was performed by the Bioinformatics Core Facility at the University of Manchester. Accession number for RNA-seq data: E-MTAB-11882 on the EMBL-EBI ArrayExpress database.

### Comet assay

A neutral comet assay was performed on day 10 cultures of mESC-derived endothelial cells treated with doxycycline or not for 24 hours to induce either TC or TC S51A expression. For a positive control, endothelial cells were treated with 30 μM H_2_O_2_ for 4 hours. The neutral comet assay was performed using the Comet SCGE Assay Kit (Enzo Lifesciences) as per manufactures instructions. Phase contrast imaging was undertaken using a Leica DMI 3000B inverted microscope fitted with an N PLAN 20x/0.35 NA PH1 objective (PH1 filter) and a DFC310FX camera. Acquisition was controlled by the Leica LAS X software.

### Immunofluorescence

Prior to immunofluorescent staining, endothelial cells were grown on 12 well chamber slides (Ibidi), coated with phenol red free matrigel (Corning) diluted to a protein concentration of 5.5 mg/ml in 1X PBS. TC or TC S51A expression was induced with dox at the indicated time points prior staining. Culture media was removed then cells were washed with 1X PBS, before fixation using 2% paraformaldehyde in 1X PBS for 20 minutes at room temperature. Cells were washed with 1X PBS, then blocking and permeabilisation buffer (1X PBS, 5% goat serum, 0.3% Triton X-100) was added to each well and incubated for 1 hour at room temperature. The blocking buffer was then removed and replaced with the primary antibodies (Supplementary Table 2) diluted in antibody dilution buffer (1X PBS, 1% BSA, 0.3% Triton X-100), and incubated for either 1 hour at room temperature or overnight at 4°C (BRCA1 antibody only). Next, each well was washed with 1X PBS, before incubation of the secondary antibodies (Supplementary Table 2) diluted in antibody dilution buffer for 1 hour at room temperature, protected from light. Cells were washed again with 1X PBS before mounting using ProLong Diamond Antifade Mounting with DAPI (Invitrogen). Slides were then left to cure overnight at room temperature, protected from light. For long term storage, slides were kept at 4°C in the dark. Imaging was carried out using the Zeiss Axio Imager M2 upright fluorescent microscope fitted with a Photometrics CoolSNAP HQ2 camera, using either a 40x/0.75 NA dry or 63x/1.3 NA oil immersion objective, as indicated. Acquisition was controlled by the Micro-Manager 2.0 software.

### Senescence associated beta-galactosidase (SA-β-gal) detection

Prior to SA-β-gal detection, mESC-derived endothelial cells were treated with dox or not for 24h prior to GFP cell sorting. Cells were then cultured for a further 4 days. The cells were then fixed for 15 minutes and stained using the Senescence Detection Kit (Abcam), as per manufacturer’s instructions. Cells were incubated in the staining solution overnight in a humidified incubator at 37°C. The plate was placed in a ziplock bag during the overnight incubation, so as the incubator CO_2_ levels did not affect colour development. Phase contrast imaging was undertaken using a Leica DMI 3000B inverted microscope fitted with an N PLAN 20x/0.35 NA PH1 objective (PH1 filter) and a DFC310FX camera. Acquisition was controlled by the Leica LAS X software.

### CRISPR/Cas9 mediated gene knockout

For deletion of *Cdkn2a*, three different crRNA sequences were used and are outlined in Supplementary Table 3 (Integrated DNA Technologies). RNA duplex was formed by combining crRNA, Alt-R tracrRNA and Duplex Buffer (all Integrated DNA Technologies) to a final concentration of 50μM, then heating at 95°C for 5 minutes before cooling to room temperature. To form the RNP complex, RNA duplex was then added to Cas9 enzyme to a final concentration of 25 μM and 24.4 μM respectively, and left at room temperature for 20 minutes. Nucleofection was performed using the Neon Transfection System (Invitrogen), as per manufacturer’s instructions. Endothelial cells were harvested and counted then resuspended in Buffer R. The RNP complex was then mixed with the cell suspension, and an electroporation protocol of 3X 10 ms pulses at 1400 V was used. Cells were then resuspended in endothelial cell media supplemented with dox or not, then re-plated at 100,000 cells/well in a 12 well plate.

### RNA imaging

Imaging of nascent RNA was performed using the Click-iT RNA Alexa Fluor 594 imaging kit (Invitrogen). Cells were incubated with 1mM 5-ethynyl uridine (5-EU) in DMSO for 1 hour prior to fixation. Staining was conducted as per manufacturer’s instructions. Imaging was carried out using the Zeiss Axio Imager M2 upright fluorescent microscope fitted with a Photometrics CoolSNAP HQ2 camera, using a 63x/1.3 NA oil immersion objective. Acquisition was controlled by the Micro-Manager 2.0 software.

## Acknowledgements

The authors thank Dr Roberto Paredes and Dr Stefan Meyer for critical reading of the manuscript. The authors thank the staff at the Flow Cytometry, Genomic Technologies, Bioinformatics and Bioimaging core facilities of the University of Manchester for technical support. This work was supported by PhD fellowship to Emily Neil by the EHE Rare Cancer Charity UK. Work in the Kouskoff laboratory is supported by the Medical Research Council (MR/P000673/1; MR/T000384/1) and the Biotechnology and Biological Sciences Research Council (BB/R007209/1).

## Author contributions

Conceptualization and methodology: EN and VK; Investigation: EN; Writing: EN and VK; Critical material provision: BR; Funding Acquisition, VK.

