## Supplemental figures for "The oncogenic fusion protein TAZ-CAMTA1 promotes genomic instability and senescence through hypertranscription"

### Slide 1
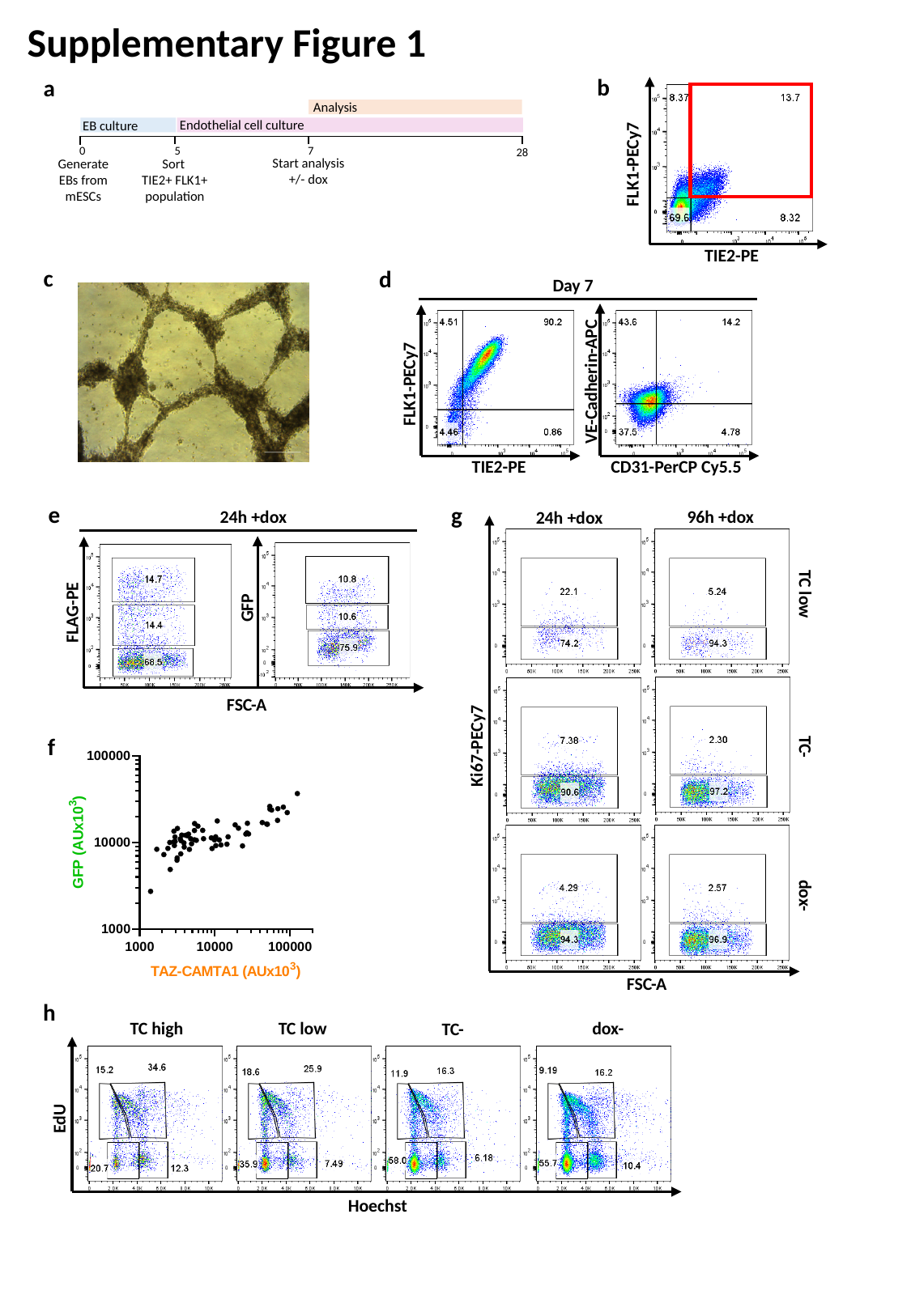

Supplementary Figure 1
b
a
FLK1-PECy7
TIE2-PE
Analysis
Endothelial cell culture
EB culture
0
7
5
28
Start analysis
+/- dox
Sort
TIE2+ FLK1+ population
Generate EBs from mESCs
c
d
Day 7
VE-Cadherin-APC
FLK1-PECy7
CD31-PerCP Cy5.5
TIE2-PE
e
g
24h +dox
96h +dox
24h +dox
Ki67-PECy7
FSC-A
GFP
FLAG-PE
FSC-A
TC low
f
TC-
dox-
h
TC high
TC low
dox-
TC-
EdU
Hoechst

### Slide 2
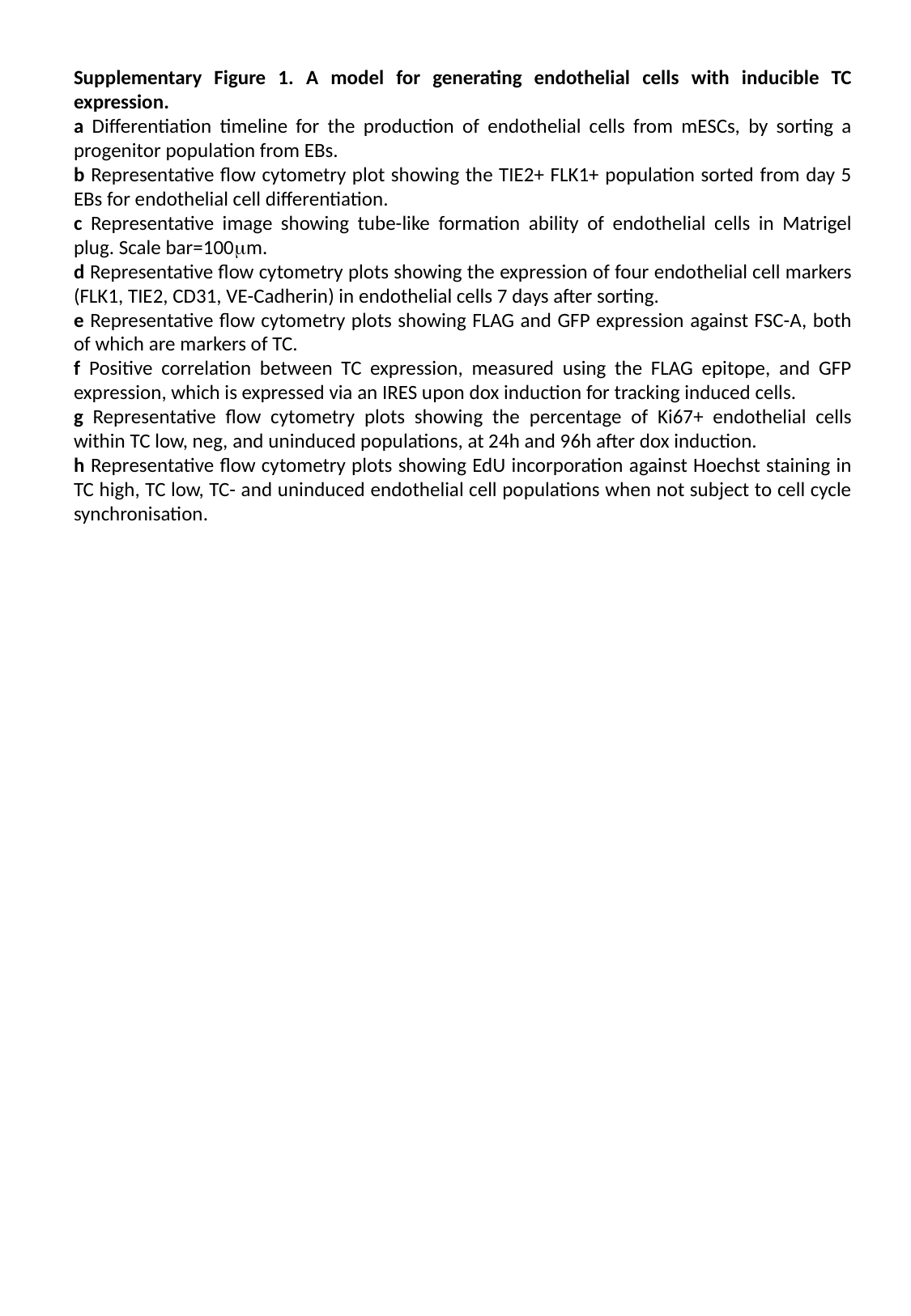

Supplementary Figure 1. A model for generating endothelial cells with inducible TC expression.
a Differentiation timeline for the production of endothelial cells from mESCs, by sorting a progenitor population from EBs.
b Representative flow cytometry plot showing the TIE2+ FLK1+ population sorted from day 5 EBs for endothelial cell differentiation.
c Representative image showing tube-like formation ability of endothelial cells in Matrigel plug. Scale bar=100μm.
d Representative flow cytometry plots showing the expression of four endothelial cell markers (FLK1, TIE2, CD31, VE-Cadherin) in endothelial cells 7 days after sorting.
e Representative flow cytometry plots showing FLAG and GFP expression against FSC-A, both of which are markers of TC.
f Positive correlation between TC expression, measured using the FLAG epitope, and GFP expression, which is expressed via an IRES upon dox induction for tracking induced cells.
g Representative flow cytometry plots showing the percentage of Ki67+ endothelial cells within TC low, neg, and uninduced populations, at 24h and 96h after dox induction.
h Representative flow cytometry plots showing EdU incorporation against Hoechst staining in TC high, TC low, TC- and uninduced endothelial cell populations when not subject to cell cycle synchronisation.

### Slide 3
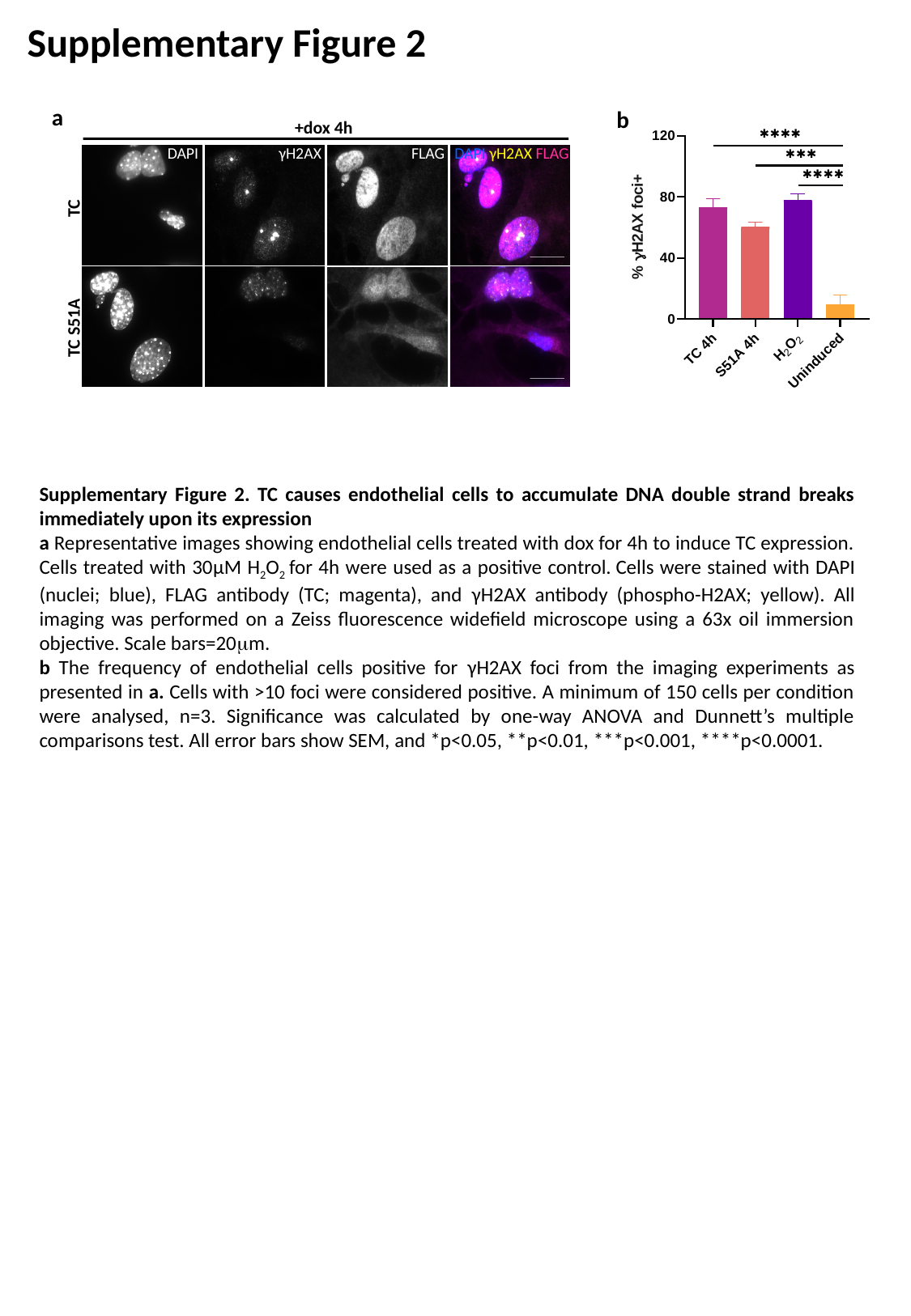

Supplementary Figure 2
a
b
+dox 4h
DAPI
γH2AX
FLAG
DAPI γH2AX FLAG
TC
TC S51A
Supplementary Figure 2. TC causes endothelial cells to accumulate DNA double strand breaks immediately upon its expression
a Representative images showing endothelial cells treated with dox for 4h to induce TC expression. Cells treated with 30μM H_2_O_2_ for 4h were used as a positive control. Cells were stained with DAPI (nuclei; blue), FLAG antibody (TC; magenta), and γH2AX antibody (phospho-H2AX; yellow). All imaging was performed on a Zeiss fluorescence widefield microscope using a 63x oil immersion objective. Scale bars=20μm.
b The frequency of endothelial cells positive for γH2AX foci from the imaging experiments as presented in a. Cells with >10 foci were considered positive. A minimum of 150 cells per condition were analysed, n=3. Significance was calculated by one-way ANOVA and Dunnett’s multiple comparisons test. All error bars show SEM, and *p<0.05, **p<0.01, ***p<0.001, ****p<0.0001.

### Slide 4
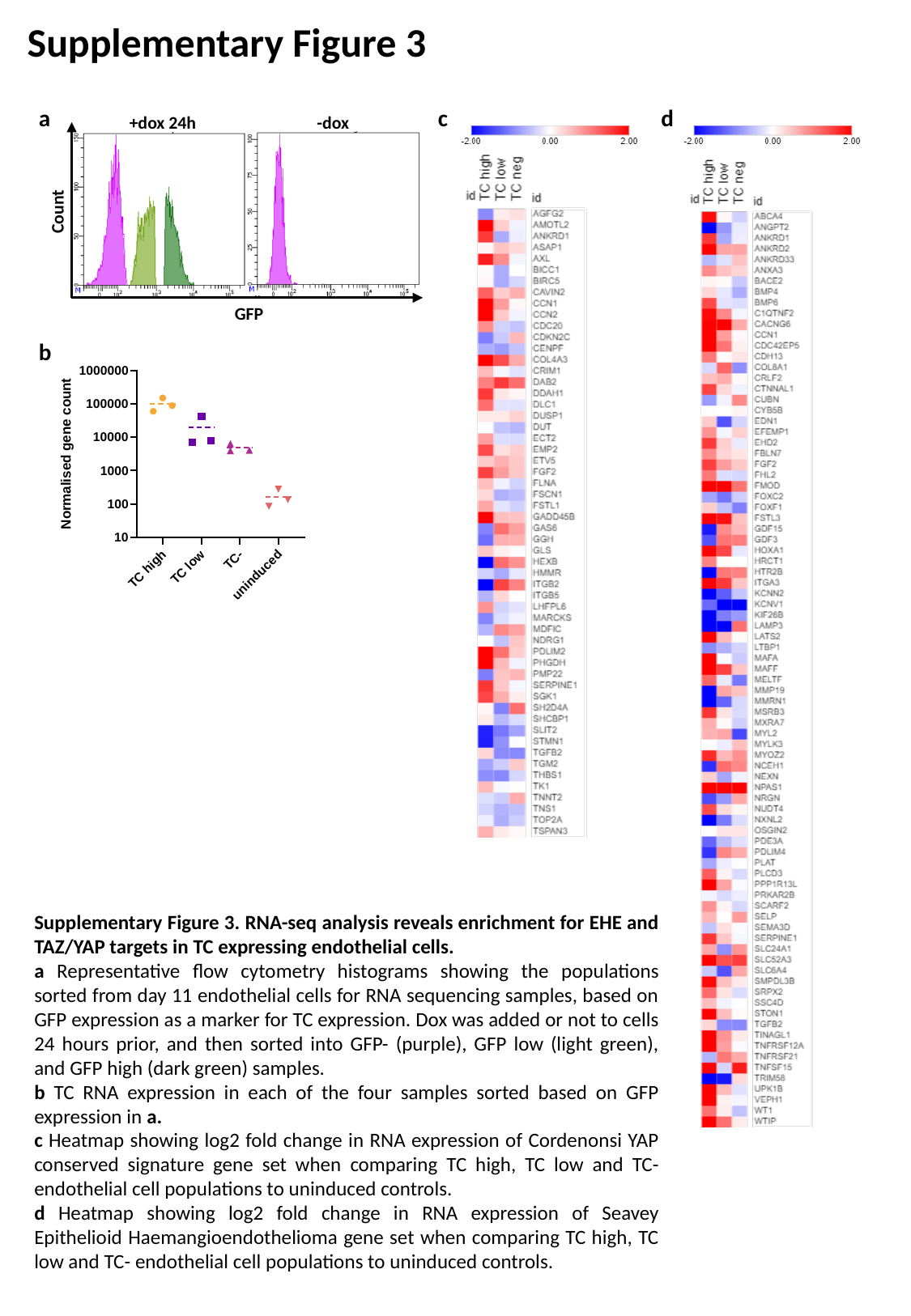

Supplementary Figure 3
d
c
a
+dox 24h
-dox
Count
GFP
b
Supplementary Figure 3. RNA-seq analysis reveals enrichment for EHE and TAZ/YAP targets in TC expressing endothelial cells.
a Representative flow cytometry histograms showing the populations sorted from day 11 endothelial cells for RNA sequencing samples, based on GFP expression as a marker for TC expression. Dox was added or not to cells 24 hours prior, and then sorted into GFP- (purple), GFP low (light green), and GFP high (dark green) samples.
b TC RNA expression in each of the four samples sorted based on GFP expression in a.
c Heatmap showing log2 fold change in RNA expression of Cordenonsi YAP conserved signature gene set when comparing TC high, TC low and TC- endothelial cell populations to uninduced controls.
d Heatmap showing log2 fold change in RNA expression of Seavey Epithelioid Haemangioendothelioma gene set when comparing TC high, TC low and TC- endothelial cell populations to uninduced controls.

### Slide 5
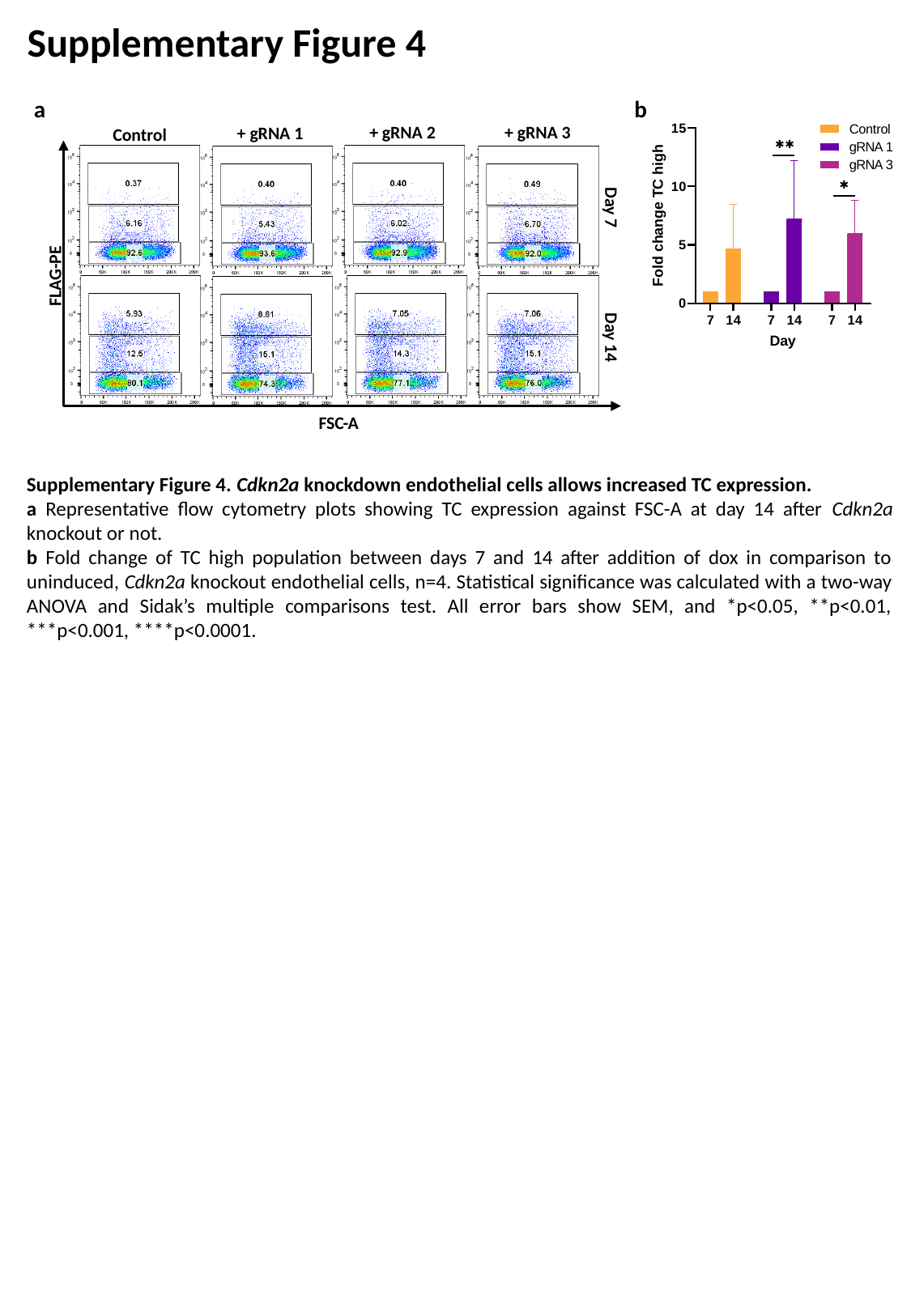

Supplementary Figure 4
a
b
+ gRNA 3
+ gRNA 2
+ gRNA 1
Control
Day 7
FLAG-PE
Day 14
FSC-A
Supplementary Figure 4. Cdkn2a knockdown endothelial cells allows increased TC expression.
a Representative flow cytometry plots showing TC expression against FSC-A at day 14 after Cdkn2a knockout or not.
b Fold change of TC high population between days 7 and 14 after addition of dox in comparison to uninduced, Cdkn2a knockout endothelial cells, n=4. Statistical significance was calculated with a two-way ANOVA and Sidak’s multiple comparisons test. All error bars show SEM, and *p<0.05, **p<0.01, ***p<0.001, ****p<0.0001.

### Slide 6
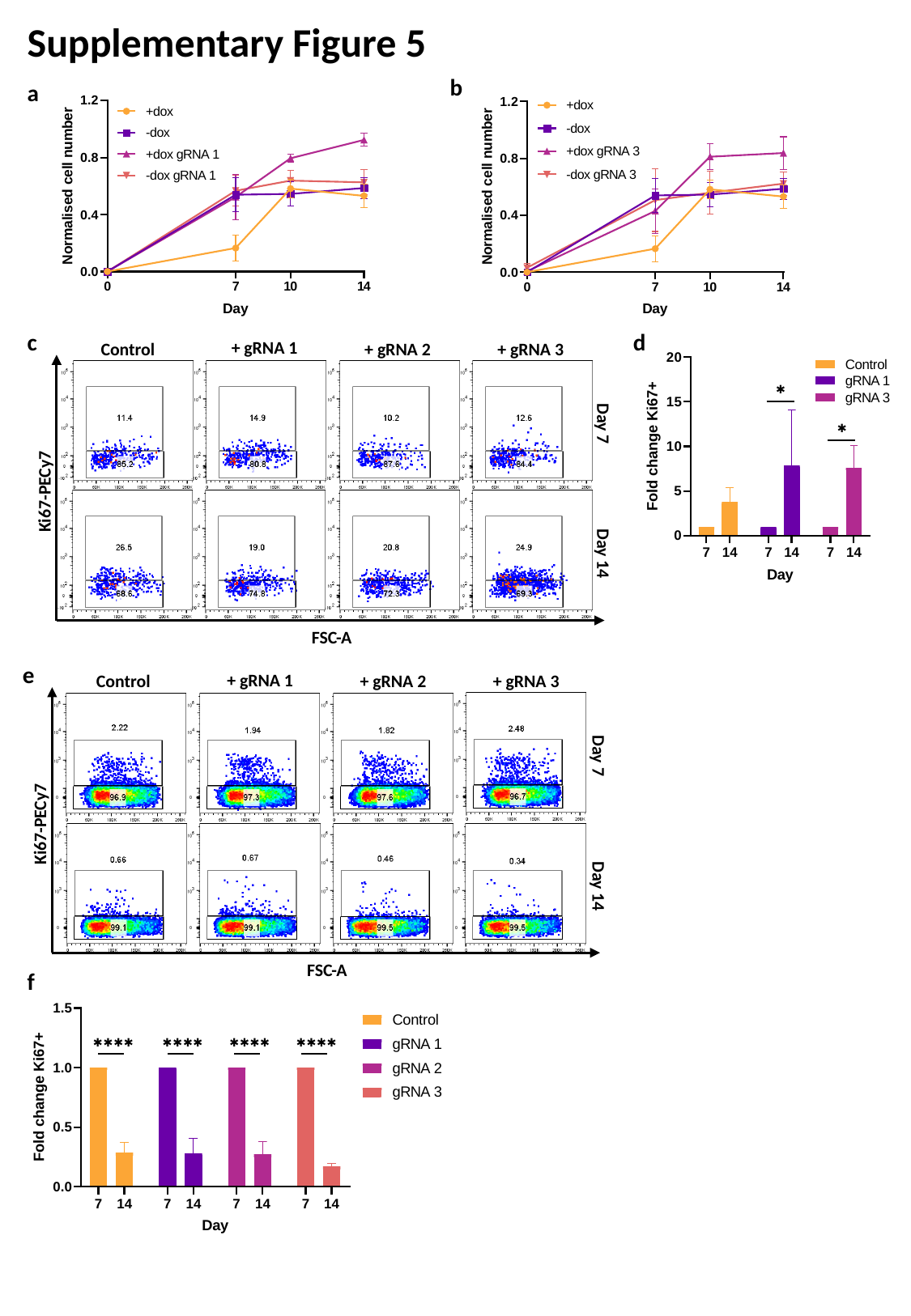

Supplementary Figure 5
b
a
d
c
+ gRNA 1
+ gRNA 3
Control
+ gRNA 2
Day 7
Ki67-PECy7
Day 14
FSC-A
e
+ gRNA 1
+ gRNA 3
Control
+ gRNA 2
Day 7
Ki67-PECy7
Day 14
FSC-A
f

### Slide 7
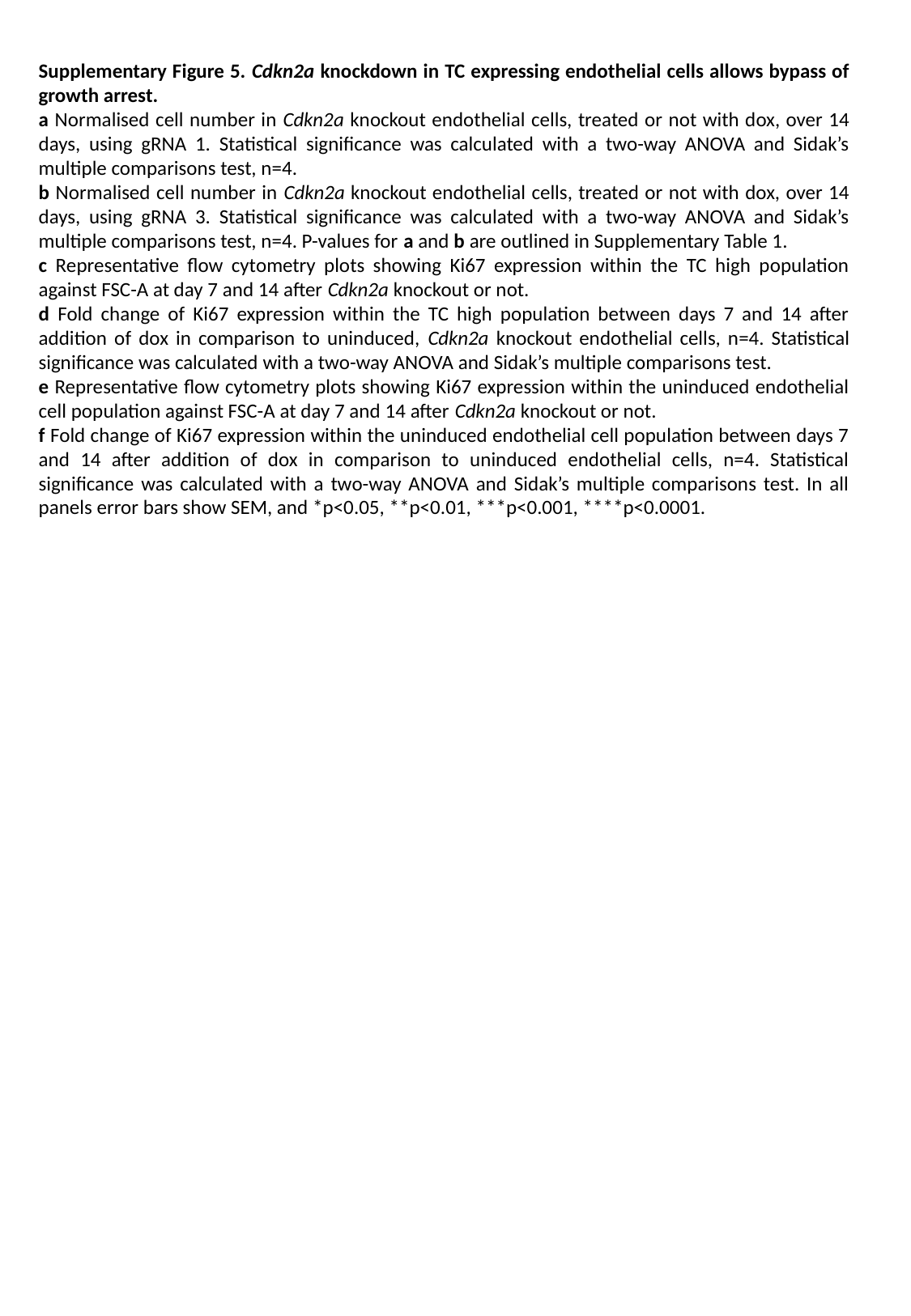

Supplementary Figure 5. Cdkn2a knockdown in TC expressing endothelial cells allows bypass of growth arrest.
a Normalised cell number in Cdkn2a knockout endothelial cells, treated or not with dox, over 14 days, using gRNA 1. Statistical significance was calculated with a two-way ANOVA and Sidak’s multiple comparisons test, n=4.
b Normalised cell number in Cdkn2a knockout endothelial cells, treated or not with dox, over 14 days, using gRNA 3. Statistical significance was calculated with a two-way ANOVA and Sidak’s multiple comparisons test, n=4. P-values for a and b are outlined in Supplementary Table 1.
c Representative flow cytometry plots showing Ki67 expression within the TC high population against FSC-A at day 7 and 14 after Cdkn2a knockout or not.
d Fold change of Ki67 expression within the TC high population between days 7 and 14 after addition of dox in comparison to uninduced, Cdkn2a knockout endothelial cells, n=4. Statistical significance was calculated with a two-way ANOVA and Sidak’s multiple comparisons test.
e Representative flow cytometry plots showing Ki67 expression within the uninduced endothelial cell population against FSC-A at day 7 and 14 after Cdkn2a knockout or not.
f Fold change of Ki67 expression within the uninduced endothelial cell population between days 7 and 14 after addition of dox in comparison to uninduced endothelial cells, n=4. Statistical significance was calculated with a two-way ANOVA and Sidak’s multiple comparisons test. In all panels error bars show SEM, and *p<0.05, **p<0.01, ***p<0.001, ****p<0.0001.

### Slide 8
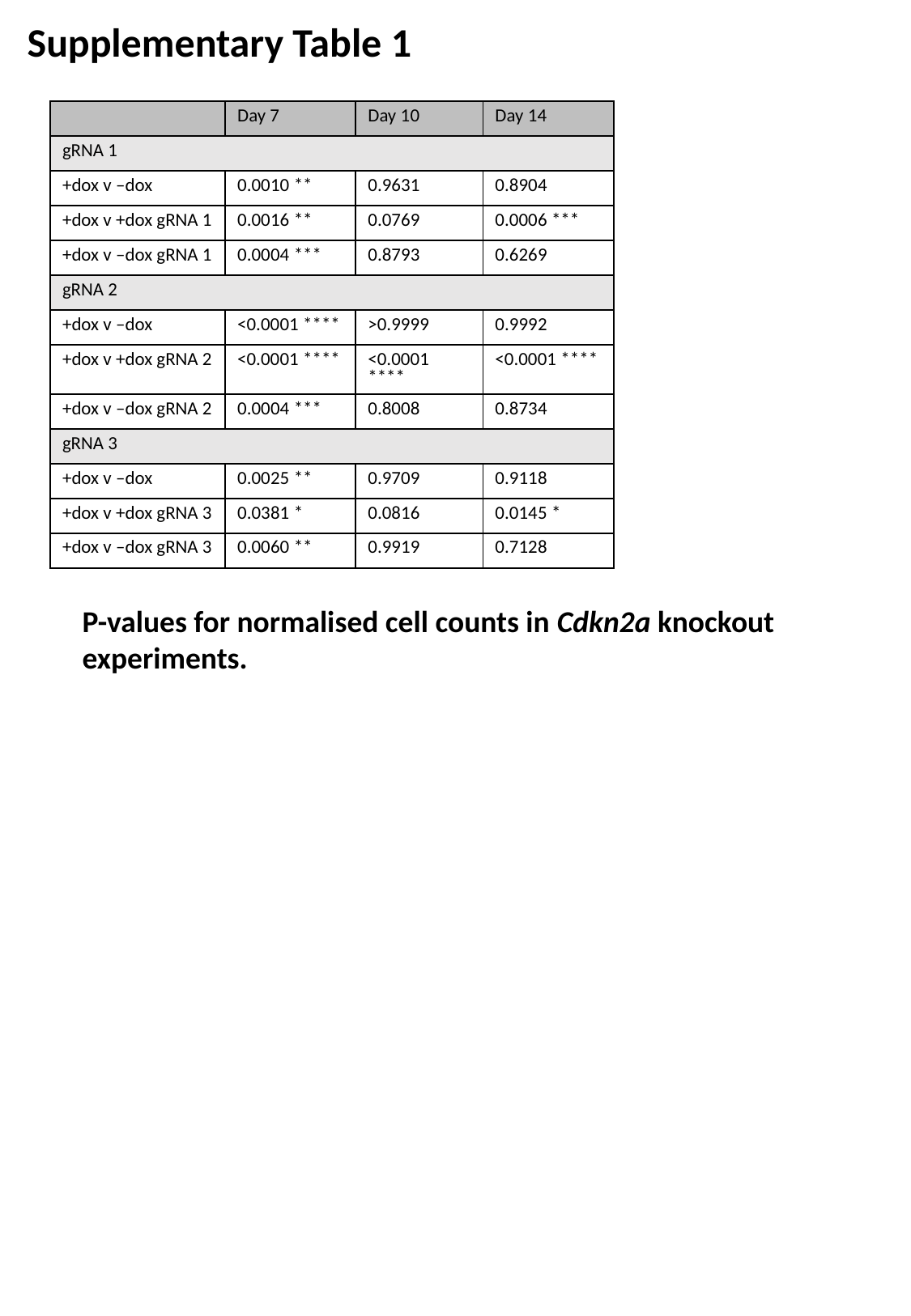

Supplementary Table 1
| | Day 7 | Day 10 | Day 14 |
| --- | --- | --- | --- |
| gRNA 1 | | | |
| +dox v –dox | 0.0010 \*\* | 0.9631 | 0.8904 |
| +dox v +dox gRNA 1 | 0.0016 \*\* | 0.0769 | 0.0006 \*\*\* |
| +dox v –dox gRNA 1 | 0.0004 \*\*\* | 0.8793 | 0.6269 |
| gRNA 2 | | | |
| +dox v –dox | <0.0001 \*\*\*\* | >0.9999 | 0.9992 |
| +dox v +dox gRNA 2 | <0.0001 \*\*\*\* | <0.0001 \*\*\*\* | <0.0001 \*\*\*\* |
| +dox v –dox gRNA 2 | 0.0004 \*\*\* | 0.8008 | 0.8734 |
| gRNA 3 | | | |
| +dox v –dox | 0.0025 \*\* | 0.9709 | 0.9118 |
| +dox v +dox gRNA 3 | 0.0381 \* | 0.0816 | 0.0145 \* |
| +dox v –dox gRNA 3 | 0.0060 \*\* | 0.9919 | 0.7128 |
P-values for normalised cell counts in Cdkn2a knockout experiments.

### Slide 9
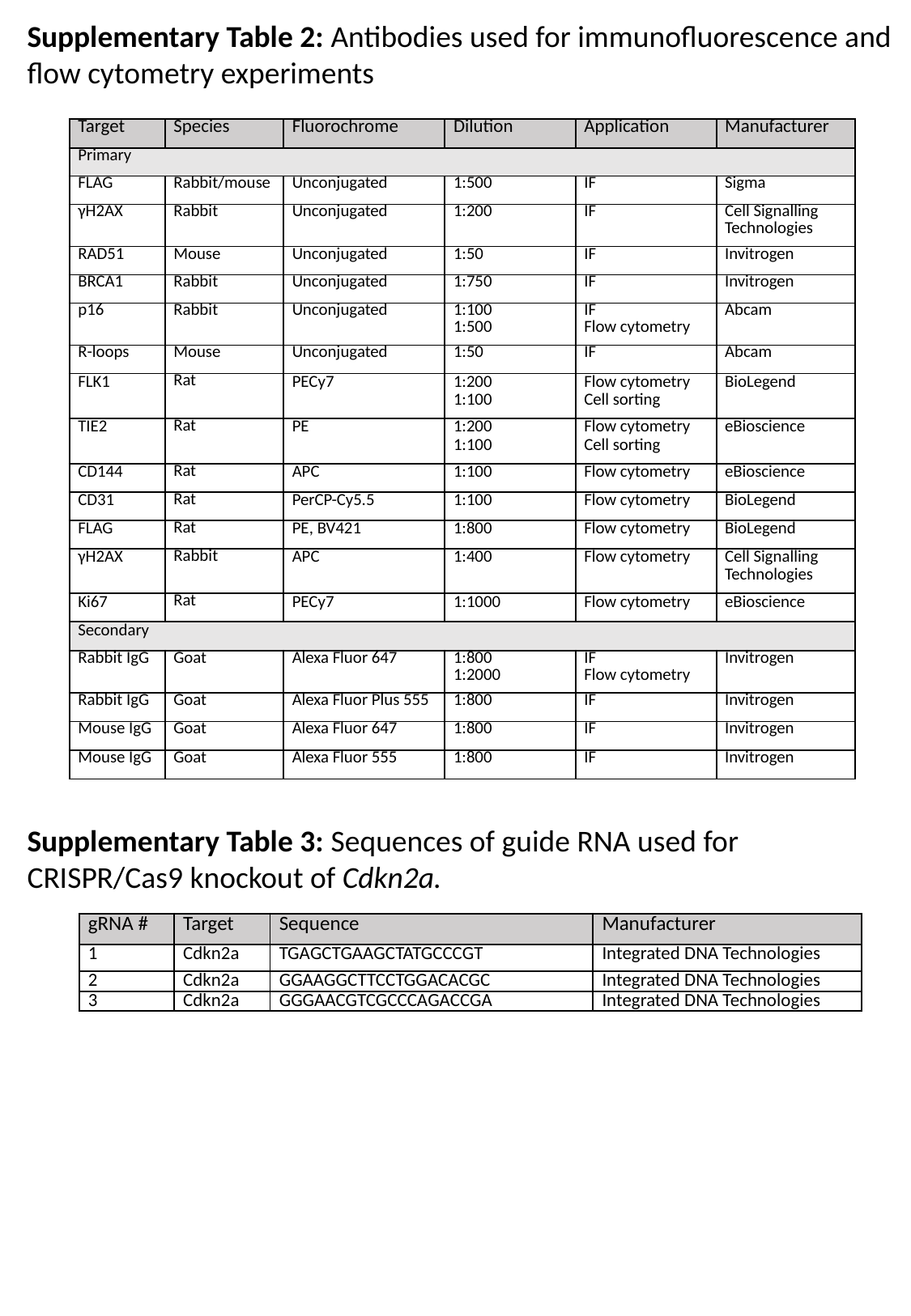

Supplementary Table 2: Antibodies used for immunofluorescence and flow cytometry experiments
| Target | Species | Fluorochrome | Dilution | Application | Manufacturer |
| --- | --- | --- | --- | --- | --- |
| Primary | | | | | |
| FLAG | Rabbit/mouse | Unconjugated | 1:500 | IF | Sigma |
| γH2AX | Rabbit | Unconjugated | 1:200 | IF | Cell Signalling Technologies |
| RAD51 | Mouse | Unconjugated | 1:50 | IF | Invitrogen |
| BRCA1 | Rabbit | Unconjugated | 1:750 | IF | Invitrogen |
| p16 | Rabbit | Unconjugated | 1:100 1:500 | IF Flow cytometry | Abcam |
| R-loops | Mouse | Unconjugated | 1:50 | IF | Abcam |
| FLK1 | Rat | PECy7 | 1:200 1:100 | Flow cytometry Cell sorting | BioLegend |
| TIE2 | Rat | PE | 1:200 1:100 | Flow cytometry Cell sorting | eBioscience |
| CD144 | Rat | APC | 1:100 | Flow cytometry | eBioscience |
| CD31 | Rat | PerCP-Cy5.5 | 1:100 | Flow cytometry | BioLegend |
| FLAG | Rat | PE, BV421 | 1:800 | Flow cytometry | BioLegend |
| γH2AX | Rabbit | APC | 1:400 | Flow cytometry | Cell Signalling Technologies |
| Ki67 | Rat | PECy7 | 1:1000 | Flow cytometry | eBioscience |
| Secondary | | | | | |
| Rabbit IgG | Goat | Alexa Fluor 647 | 1:800 1:2000 | IF Flow cytometry | Invitrogen |
| Rabbit IgG | Goat | Alexa Fluor Plus 555 | 1:800 | IF | Invitrogen |
| Mouse IgG | Goat | Alexa Fluor 647 | 1:800 | IF | Invitrogen |
| Mouse IgG | Goat | Alexa Fluor 555 | 1:800 | IF | Invitrogen |
Supplementary Table 3: Sequences of guide RNA used for CRISPR/Cas9 knockout of Cdkn2a.
| gRNA # | Target | Sequence | Manufacturer |
| --- | --- | --- | --- |
| 1 | Cdkn2a | TGAGCTGAAGCTATGCCCGT | Integrated DNA Technologies |
| 2 | Cdkn2a | GGAAGGCTTCCTGGACACGC | Integrated DNA Technologies |
| 3 | Cdkn2a | GGGAACGTCGCCCAGACCGA | Integrated DNA Technologies |
